# A consensus genome sequence for the social amoeba *Dictyostelium giganteum*

**DOI:** 10.64898/2026.03.01.708701

**Authors:** Aditya Sharma, Kumari Khushi, Febina Ravindran, Jayarama S. Kadandale, Bibha Choudhary, Subha Srinivasan, Vidyanand Nanjundiah

## Abstract

The life cycle of the Dictyostelid amoebae is unusual in that it alternates between a free-living solitary phase and an aggregative social phase. We used six previously collected *Dictyostelium giganteum* strains from distinct ecological niches in the Mudumalai Nature Reserve, India. From them, we generated short Illumina reads and assembled a consensus genome, comprising nuclear and mitochondrial genomes, representative of all six strains. The nuclear assembly has an AT content of 75.76%, accounting for 38.52 Mb, and resolves into five chromosome-scale scaffolds that are consistent with the published karyotype. Its N50 is 3.01 Mb and L50 is 5. The BUSCO analysis shows 3.9% fragmentation and 92.1% completeness. Genome assembly and completeness are also validated using ∼5,500 genes from a publicly available transcriptome dataset (PRJNA48443) derived from the post-aggregation stage. 13,251 predicted proteins are encoded by the genome, including ABC transporters, polyketide synthases, Ras/Rho GTPases, and expanded families of protein kinases. Comparative analysis demonstrates extensive conservation of syntenic blocks related to the dictyostelids *D. discoideum* and *D. firmibasis* as well as lineage-specific rearrangements. About 18% of the nuclear genome is made up of repetitive DNA, mostly in the form of simple repeats. Major transposable element classes, including piggyBac-like fragments, were found by homology searches. Long poly-asparagine/glutamine tracts are less common than in *D. discoideum*, but low-complexity sequences are common due to strong AT-driven codon bias. Comparative proteome-level orthology analysis across Dictyostelium species and Entamoeba identified a conserved Amoebozoan core together with a substantial Dictyostelium-specific gene repertoire. Domain-level comparisons further revealed widespread conservation of intracellular signalling and cytoskeletal modules shared with animals, whereas canonical metazoan extracellular adhesion domains were absent, highlighting the deep evolutionary roots of regulatory complexity underlying aggregative multicellularity.

## INTRODUCTION

The generation of whole genome sequences has expanded the range of questions that can be asked regarding the development, evolution and ecology of organisms. Besides, it has opened the prospect of their better utilisation for attacking general problems in diverse areas of biology, including, lately, human disease. Whole genome sequencing enables one to ask which features of an organism’s genome are shared with others and which are special to it, following which one can attempt to correlate those features with phenotypic traits and conjecture evolutionary explanations. The combination of diversity and similarity among the Cellular Slime Moulds (CSMs) makes them especially attractive objects of study from that viewpoint.

CSMs are Eukaryotes belonging to the group Mycetozoa within the super-group Amoebozoa. They are ubiquitous in soils all over the world (Raper 2014) and are also found in animal droppings (Brefeld 1884; Stephenson and Landolt 1992; Suthers 1985). They occur all over the world; dispersion of spores by animal vectors, in particular birds, is a plausible reason for their cosmopolitan distribution. They have also been reported from two unexpected environments: a human cornea, in which *D. polycephalum* (renamed *Coremiostelium polycephalum*) was associated with keratitis (Reddy et al. 2010), and ascocarps of an edible mushroom (Hu et al. 2024). The CSMs are paradigms for the evolution of aggregative multicellularity (Niklas and Newman 2016). They serve to illustrate evolutionary convergence, not only across genera (Bonner 2009), but also between eubacteria and eukaryotes (Arias Del Angel et al. 2020). Further, they are excellent models for cell, developmental, and disease biology (Escalante and Cardenal-Muñoz 2019). Under the stress of starvation, groups of free-living CSM amoebae aggregate via chemotaxis and form a sorocarp or fruiting body. Such sorocarp formation is a convergent trait seen in five of the six major eukaryotic supergroups (Brown and Silberman 2013). CSMs display convergence of another sort. Traits that were believed at one time to be species-specific show up sometimes - albeit rarely - in other species (Bonner 2003).

High-quality genome assemblies of *Dictyostelium discoideum* and *Dictyostelium firmibasis*, publicly available through dictyBase (Fey et al. 2009; Basu et al. 2015), have provided foundational genomic references for Dictyostelia. In the present study, these resources were used for comparative analyses, reference-guided scaffolding, and evolutionary interpretation of the *Dictyostelium giganteum* genome.

Importantly, the *D. giganteum* genome presented here was assembled exclusively from short-read Illumina sequencing data. Despite the inherent challenges of reconstructing an AT-rich and repeat-dense genome using short reads alone, we achieved a near-complete assembly resolving into five chromosome-scale scaffolds consistent with the known karyotype. High sequencing depth, stringent contamination filtering, iterative polishing, reference-guided scaffolding, and progressive multi-strain integration collectively enabled high contiguity and completeness.

Here, we present the nuclear and mitochondrial genome sequences of the cellular slime mold *D. giganteum* (Singh 1947) and analyze them with respect to genome content, structural organization, functional composition, and phylogenetic context. The remainder of the paper describes the strains used and the experimental and computational strategies employed, followed by a comprehensive characterization of genome architecture and comparative analyses with *D. discoideum* and *D. firmibasis*. By generating a near-complete, chromosome-scale assembly derived from multiple wild strains and integrating comparative, functional, and evolutionary analyses, this study establishes *D. giganteum* as a well-resolved genomic model within Dictyostelia. The resulting genomic resource provides a robust framework for investigating genome evolution, developmental regulation, natural variation, and the genetic basis of aggregative multicellularity.

## MATERIALS AND METHODS

### Sample Collection and Preparation

#### Source and strain descriptions

Six Dictyostelium giganteum strains were obtained from two ecological niches representing distinct ecological niches respectively within and adjoining a 50-hectare undisturbed forest plot the Mudumalai Nature Reserve, India (Kaushik et al. 2006; Sathe et al. 2010). Strains 46a3 and 46c6 were isolated as soil-dwelling organisms, while strains F4, F5, F15, and F16 were derived from distinct spores of a single fruiting body collected from elephant dung within the same reserve. This sampling approach enabled analysis of both environmental diversity and potential clonal variation within a single fruiting body.

#### DNA extraction protocols

Genomic DNA from *Dictyostelium giganteum* strains was extracted via standard phenol-chloroform protocol, lysing amoebae in SDS buffer with proteinase K, followed by phenol: chloroform: isoamyl alcohol (25:24:1) extraction, ethanol precipitation with sodium acetate at –20°C, 70% ethanol wash, and resuspension in TE buffer.

#### Library construction and Sequencing

Sequencing libraries were prepared using the NEBNext® Ultra™ II DNA Library Prep Kit according to the manufacturer’s instructions. For physically sheared DNA, 500 pg of input DNA was used per reaction. DNA fragmentation was performed either enzymatically (using the NEBNext Ultra II FS kit) or by mechanical shearing. Fragmented DNA underwent end repair and A-tailing to generate 3′ A-overhangs. Adapters were then ligated to the repaired DNA fragments, followed by PCR enrichment to amplify adaptor-ligated molecules. Libraries were purified and size-selected using SPRIselect magnetic beads.

All six strains were sequenced on the Illumina HiSeq 2500 platform to generate paired-end reads suitable for de novo genome assembly and comparative genomic analyses.

### Sequencing Data Processing

#### Adapter removal

Adapters were trimmed using Trimmomatic v0.39, and high-quality reads were extracted for downstream analysis (Bolger et al. 2014).

#### Quality control

The quality of raw reads was evaluated using FastQC, providing insights into sequencing quality, adapter contamination, and overall read characteristics (https://www.bioinformatics.babraham.ac.uk/projects/fastqc/)

#### Contamination removal

To identify and classify this contamination, Kraken2 v2.0.9-beta was employed using a k-mer-based approach for rapid and accurate taxonomic classification of sequencing reads (Wood et al. 2019). A pre-built 4 GB MiniKraken database, containing complete bacterial, archaeal, and viral genomes from RefSeq (as of October 18, 2017), was used for classification. After classifying the Illumina reads, the Kraken2-generated taxonomic reports were visualized using matplotlib (Python 3.10), enabling graphical representation of the contamination profile in the dataset.

### Genome Assembly

#### Contig-level

High-quality, adapter- and quality-trimmed reads from all six strains were assembled de novo using **MEGAHIT** (Li et al. 2015) with default parameters optimized for complex eukaryotic genomes.

#### Scaffolding

Contig assemblies were scaffolded using **Ragtag** (Alonge et al. 2022) in reference-guided mode, with the chromosome-level assembly of *Dictyostelium firmibasis* as the reference genome.

#### Polishing

Final polishing was performed with Pilon to correct base-level errors and enhance overall assembly accuracy (Walker et al. 2014).

### Assessment of Assemblies

Assembly quality and completeness were assessed using QUAST v5.3.0, which provided assembly statistics such as total length, contig and scaffold N50/L50 values, and misassembly rates (Gurevich et al. 2013). Additionally, BUSCO v5.5.0 analysis was conducted to evaluate genome completeness based on conserved orthologous gene sets (Manni et al. 2021).

### Constructing a Consensus Genome

A species-level consensus reference genome for *Dictyostelium giganteum* was generated through a progressive, alignment-based integration of assemblies from six independently sequenced strains. Individual assemblies were first indexed using SAMtools v1.18 (Li et al. 2009), and assembly statistics such as total genome size, contig count, and N50 were calculated to document baseline characteristics. The assembly of strain 46a3 was selected as the initial reference backbone for integration. Additional strain assemblies were successively aligned to the backbone using NUCmer from MUMmer v4.0.0rc1 (Marçais et al. 2018). Alignment statistics were calculated and aligned regions exhibiting ≥90% sequence identity and consistent synteny were selected for incorporation.

Consensus construction was performed iteratively: aligned regions from each strain assembly were compared with the current reference, and matching segments were integrated to extend or correct corresponding regions while preserving the structural integrity of the backbone. This process was repeated for each strain until no further high-confidence additions could be made.

### Mitochondrial Genome Analysis

The mitochondrial genome of *Dictyostelium giganteum* was identified from the whole-genome assembly using BLAST searches against mitochondrial genomes of *D. discoideum* and *D. firmibasis*. Protein-coding genes were predicted by ORF detection and annotated through similarity searches against curated mitochondrial datasets and the NCBI nr database.

### Comparative Genomics

Comparative genomic analyses were conducted to assess inter-species genome organization in *Dictyostelium giganteum*. Whole-genome alignments were performed using NUCmer (from MUMmer v4.0.0rc1) to identify conserved and rearranged genomic regions across assemblies. Syntenic relationships and large-scale structural variations were visualized using a **python script**, enabling the identification of collinear blocks, inversions, and translocations between related *Dictyostelium* species.

### Repeat Element Identification

Simple sequence repeats (SSRs) were identified and characterized using MISA, enabling the detection of microsatellite distributions across the assembled genomes (Thiel et al. 2003). Transposable elements (TEs) were detected using tBLASTx v2.14.0 (Camacho et al. 2009) with an e-value threshold of 1×10⁻¹⁵ against reference TE sequences obtained from Repbase (Genetic Information Research Institute). To ensure taxonomic relevance, the Repbase database was filtered for *Dictyostelium discoideum* elements prior to analysis (Bao et al. 2015).

### Transcriptome Data Processing and Assembly

RNA-seq data for *Dictyostelium giganteum* were retrieved from the NCBI Sequence Read Archive (BioProject PRJNA48443). Raw reads were quality-checked and adapter-trimmed prior to de novo transcriptome assembly using Trinity (Grabherr et al. 2011). The resulting transcript models were aligned to the *D. giganteum* genome using GMAP to obtain splice-aware mappings and assess transcript support for annotated genes (Wu and Watanabe 2005). Functional annotation of assembled transcripts was performed using Swiss-Prot, and stage-specific expression patterns were inferred by comparison with established developmental markers from *Dictyostelium discoideum*.

### Non-Coding RNA Annotation

Transfer RNA (tRNA) genes were predicted using tRNAscan-SE based on conserved sequence and structural features (Chan et al. 2021). Other classes of non-coding RNAs (ncRNAs), including rRNAs, spliceosomal snRNAs, and snoRNAs, were identified using Infernal v1.1.4 with Rfam covariance models restricted to Amoebozoa (Nawrocki and Eddy 2013; Kalvari et al. 2021).

Ribosomal DNA (rDNA) loci were identified through sequence similarity searches and repeat analysis, and their genomic organization was examined to assess potential extrachromosomal and palindromic structures. Comparative analyses were performed against *Dictyostelium discoideum* and *Dictyostelium firmibasis* to evaluate conservation of rDNA architecture and ncRNA repertoires. Motif searches were conducted to detect Dictyostelium upstream sequence elements (DUSE).

### Gene Prediction

Gene prediction was conducted using AUGUSTUS v3.2.3, a widely used tool for eukaryotic genome annotation using publicly available genome, transcriptome, and annotation resources from *Dictyostelium discoideum*. This custom-trained model was then applied to our assembled genomes to enhance the accuracy of exon–intron structure prediction and overall gene annotation quality. The resulting high-confidence gene models were retained for downstream analyses (Stanke et al. 2004).

### Functional Annotation

Predicted protein sequences were functionally annotated using eggNOG-mapper to assign COG categories, Gene Ontology terms, KEGG pathways, and orthology-based functional predictions (Huerta-Cepas et al. 2017). Protein domain and motif annotations were obtained using InterProScan (Jones et al. 2014), integrating signatures from multiple databases, including Pfam. Additional similarity-based functional evidence was obtained through BLASTp searches against the NCBI non-redundant (nr) protein database (Altschul et al. 1990; NCBI Resource Coordinators et al. 2018). Gene family clustering and orthogroup inference across *Dictyostelium giganteum*, *D. discoideum*, *D. firmibasis*, and *Entamoeba* were performed using orthology-based clustering approaches to identify conserved and lineage-specific gene families. Domain abundance and family expansion analyses were derived from Pfam and orthogroup classifications.

### Comparative Analysis of the Metazoan Developmental Toolkit

To assess whether elements of the canonical “metazoan developmental toolkit” predate animals, we performed a domain-level comparison across Amoebozoa and Metazoa. The toolkit was defined as a curated set of Pfam domains associated with animal adhesion, developmental signalling, transcriptional regulation, apoptosis, and cytoskeletal organization, based on prior studies of early animal evolution (Newman and Bhat 2008; Brunet and King 2017; Xiao et al. 2025).

Proteomes from *Entamoeba*, three Dictyostelium species, and representative metazoans were annotated with InterProScan v5. Pfam domains were scored as present if detected in at least one protein per lineage. Domain distributions and copy numbers were used to infer conservation and lineage-specific expansion without assuming strict gene orthology.

### Horizontal Gene Transfer and Disease Gene Analysis

Putative horizontal gene transfer (HGT) candidates were identified by screening Pfam domain annotations for bacterial-associated domains and evaluating taxonomic distribution using BLASTp searches against the NCBI non-redundant database. Presence–absence comparisons were performed against *Dictyostelium discoideum* and related dictyostelids to assess conservation patterns.

Human disease-associated orthologues were identified by querying a curated set of human disease proteins against the *D. giganteum* predicted proteome using BLASTp with e-value ≤ 1e-10 cutoff. High-confidence matches were retained based on sequence similarity and alignment coverage.

## RESULTS

### Nuclear Genome Assembly

The consensus genome of *Dictyostelium giganteum* spans 38.52 Mb and is composed of 3,559 contigs, with an N50 length of 3.01 Mb and an L50 of 5. The assembly was generated at approximately 340× coverage. A total of 0.81 Mb (2.11%) consists of undetermined bases distributed across 2,652 gaps longer than 10 bp.

In addition to overall assembly metrics, compositional features were examined. The genome is AT-rich (75.76%) and exhibits broadly uniform base composition at the whole-genome scale. Genome-wide analysis revealed depletion of CpG dinucleotides with a corresponding enrichment of TpG dinucleotides, consistent with previously reported compositional features of Dictyostelium genomes (Eichinger et al., 2005).

### Chromosome Assignment

Following assembly characterization, chromosomal structure was evaluated. The nuclear genome assembly resolves into five chromosome-equivalent scaffolds (Figure 1), which is in accordance with the karyotype (Kumar et al. 2024). Following established convention, and also as has been done in the case of *D. firmibasis* (Edelbroek et al. 2024), Chromosomes are numbered in decreasing order of scaffold size, thereby providing a clear and reproducible framework that aligns with the physical karyotype. The mitochondrial genome and ribosomal DNA emerge as separate, distinct contigs that are not part of the chromosome scaffolds.

**Figure 1:**
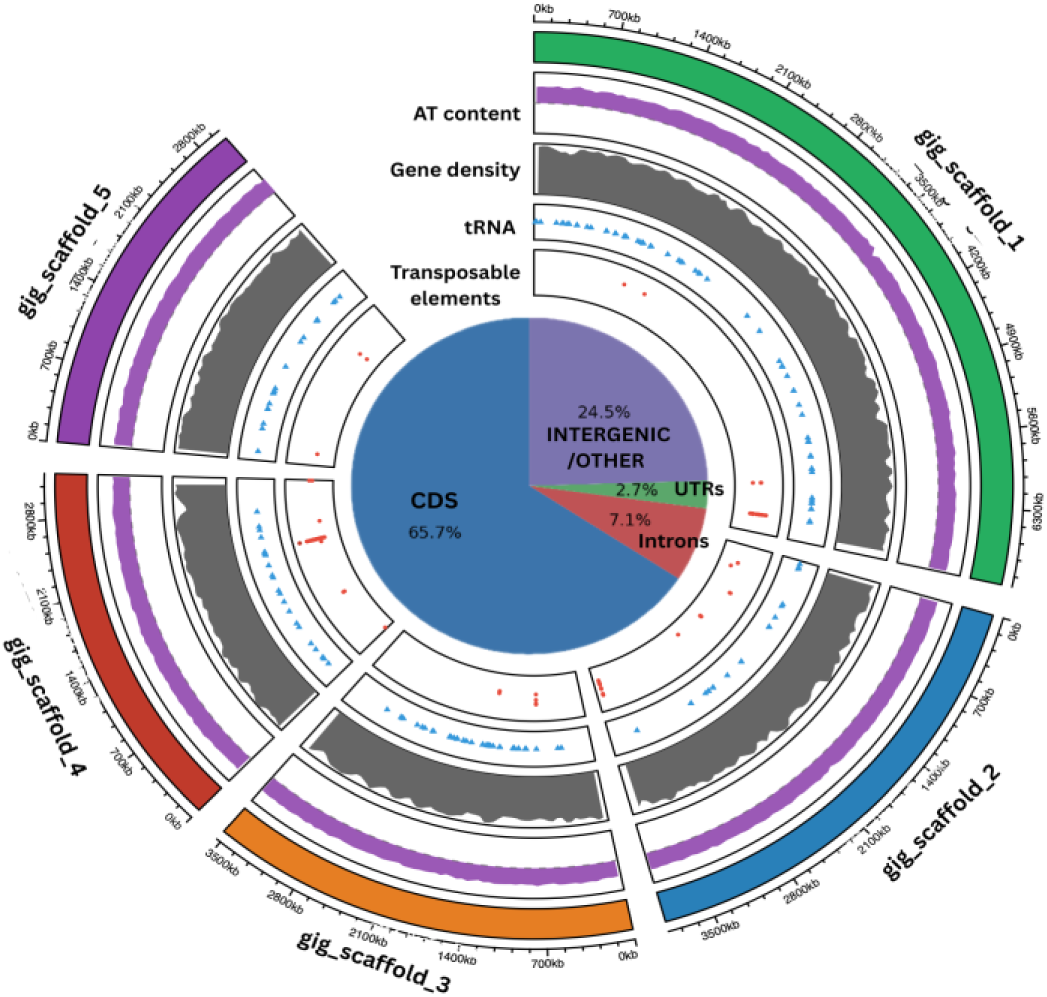
Circular representation of the *Dictyostelium giganteum* genome assembly and annotation: Figure 1 is a circos pictorial representation of the consensus genome of *Dictyostelium giganteum*. It provides an overview of genome structure, gene organization, and repetitive element distribution across the genome. Each of the five outer segments represents a major assembled scaffold. Going inwards from the outermost circle, (i) assembled scaffolds are represented in distinct colours (with genomic coordinates in kilobases); (ii) AT content is shown as a continuous purple track; (iii) gene density is in the form of a grey histogram; (iv) predicted tRNA loci are indicated in blue triangles; and (v) transposable element locations are marked by red dots. The central pie chart shows the proportion of the genome made up of potential protein-coding sequences (CDS, 65.7%), introns (7.1%), untranslated regions (UTRs; 2.7%), and regions with intergenic plus possibly regulatory sequences (24.5%).

### Comparative synteny analysis

To place the assembly in an evolutionary context, whole-genome synteny analysis was performed. Whole-genome synteny analysis reveals extensive conservation between *Dictyostelium giganteum* and both *D. firmibasis* and *D. discoideum* (Figure 2). Clear syntenic relationships are observed between the five largest *D. giganteum* scaffolds, which correspond to the five nuclear chromosomes, and chromosomes of the two reference species. These relationships include both one-to-many and many-to-one correspondences, indicating the presence of conserved chromosomal blocks alongside chromosomal rearrangements among the three species.

**Figure 2:**
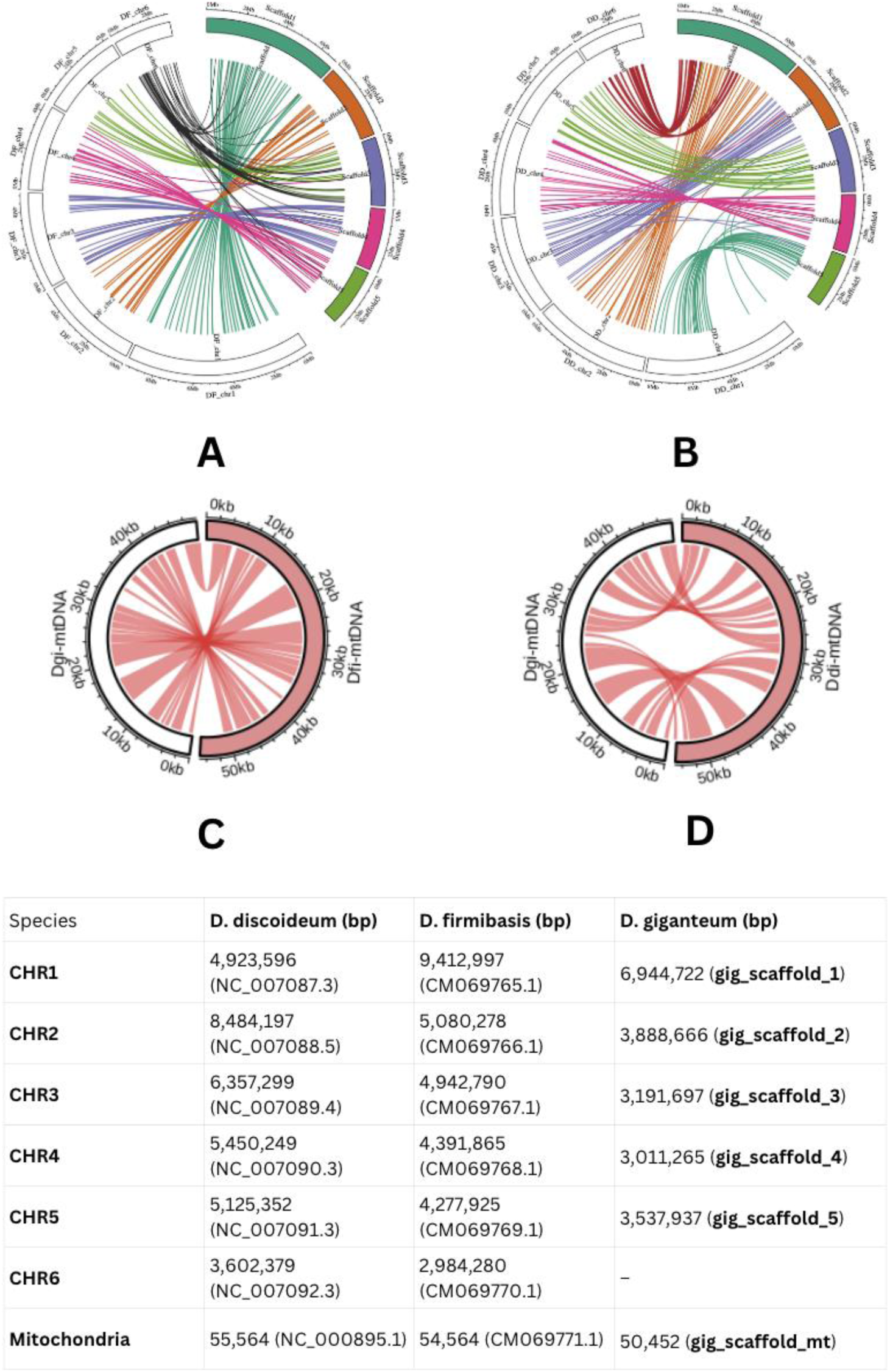
Comparative nuclear and mitochondrial synteny between *Dictyostelium giganteum*, *D. discoideum*, and *D. firmibasis*: Circular synteny plots illustrate genome-wide conservation between *Dictyostelium giganteum* and related species. (A) Nuclear genome synteny between the five largest *D. giganteum* scaffolds (Scaffold1–5; colored outer segments) and chromosomes of *D. firmibasis* (inner tracks). (B) Nuclear genome synteny between the same *D. giganteum* scaffolds and chromosomes of *D. discoideum*. In panels A and B, colored ribbons represent homologous genomic blocks identified by BLASTn alignments, highlighting extensive conservation of large syntenic segments alongside lineage-specific rearrangements and scaffold–chromosome reassignments. (C) Mitochondrial genome synteny between *D. giganteum* and *D. firmibasis*. (D) Mitochondrial genome synteny between *D. giganteum* and *D. discoideum*.

At the mitochondrial level, circular synteny plots (Figure 2) illustrate conservation of gene content and partial conservation of gene order among the mitochondrial genomes of *D. giganteum, D. firmibasis,* and *D. discoideum*, with localized rearrangements.

### Mitochondrial Genome Assembly

In parallel with nuclear genome assembly, mitochondrial sequencing was conducted. Sequencing and assembly of *Dictyostelium giganteum* produced a single mitochondrial scaffold of 50,452 bp, representing a complete mitochondrial genome which is also AT-rich (75.7% AT). The mitochondrial genome of *Dictyostelium giganteum* encodes 36 protein-coding genes, comprising core components of the mitochondrial oxidative phosphorylation machinery and the mitochondrial translation system. Coding sequences exhibit a pronounced AT-biased codon usage, with TTA (Leu) and AAA (Lys) representing the most frequently used codons, consistent with the overall AT-rich nucleotide composition of the genome.

Mitochondrial gene organization is highly polarized, with all open reading frames encoded on a single strand. Annotation of non-coding RNA elements identified a conserved set of 14 mitochondrial tRNA amino acid types, together with a complete small-subunit rRNA (rns, 12S) and fragmented large-subunit rRNA loci (rnl, 16S). In addition, multiple mitochondrial ribosomal protein genes (rpl and rps) were identified, constituting the core components of the mitochondrial translation apparatus.

For panels C and D, circular plots represent complete mitochondrial genomes scaled by length, with tick marks indicating genomic coordinates (kb). Red ribbons denote homologous sequence blocks, revealing overall conservation of mitochondrial gene order with localized structural rearrangements.

The accompanying table summarizes chromosome- and scaffold-level sizes and accession identifiers for *D. discoideum*, *D. firmibasis*, and *D. giganteum*, including mitochondrial genome lengths, providing the genomic context for the synteny relationships depicted in panels A–D.

### Repetitive and other noncoding DNA

Following the structural characterization of the genome, repetitive content was examined. 5,248,827 bp, or ∼18%, of the *D. giganteum* nuclear genome is made up of repetitive DNA of various types. The major repeat classes and their sizes are shown in Table 1.

**Table 1:** Summary of repeat classes showing the number of elements, total length (bp), and percentage of genome occupied by each repeat category.

| Repeat class | Count | Total length (bp) | % of genome |
| --- | --- | --- | --- |
| Simple repeats | 86,157 | 4,057,716 | 14.1 |
| Unknown repeats | 5,526 | 1,105,987 | 3.9 |
| Unspecified repeats | 579 | 71,206 | 0.25 |

#### Transposable Elements

To further characterize repetitive content, transposable elements (TEs) were analyzed in detail. Transposable elements (TEs) in *Dictyostelium giganteum* were identified by tblastx searches against the Repbase database, revealing representatives from all major TE classes—DNA transposons, LTR retrotransposons, and non-LTR retrotransposons (Supplementary TableS1). Among the DNA transposons, families such as DINOLT, DRE, and piggyBac were detected (not reported in D. discoideum), each represented by a limited number of short, fragmented copies. The LTR retrotransposons include DGLT-A, Gypsy, and DIRS1 families. Notably, DIRS1 shows extensive sequence conservation and the highest number of homologous fragments, indicating that it remains one of the most structurally preserved and possibly historically active LTR elements in *D. giganteum*. The non-LTR retrotransposons, comprising the TRE3 and TDD families, are diverse and form the largest TE category.

**Figure 3A:**
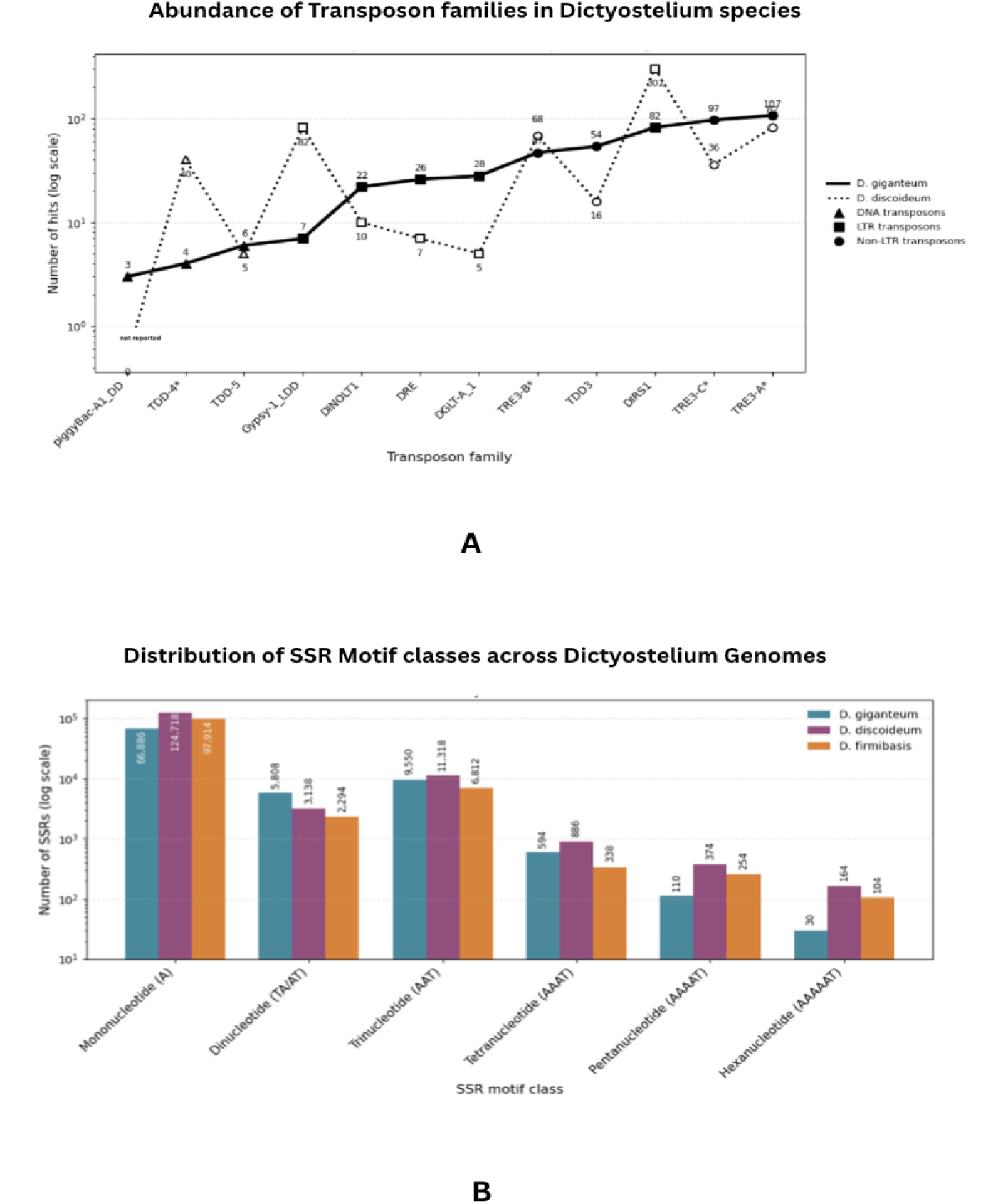
Abundance of transposon families in *Dictyostelium* genomes. **Figure 3B: Distribution of simple sequence repeat (SSR) motif classes across *Dictyostelium* genomes. (A)** Line plot showing the copy number of representative transposon families in *Dictyostelium giganteum* and *D. discoideum*. The x-axis lists transposon families ordered as plotted, and the y-axis indicates the number of hits on a logarithmic scale. Solid lines represent *D. giganteum*, whereas dotted lines represent *D. discoideum*. Marker shapes denote transposon class: triangles indicate DNA transposons, squares indicate LTR retrotransposons, and circles indicate non-LTR retrotransposons. Numeric labels indicate family-specific copy numbers. For TRE3 elements, *D. discoideum* labels are positioned above the dotted line to avoid overlap with *D. giganteum*. **(B)**Bar plot showing the abundance of SSRs classified by motif length (mono- to hexanucleotide repeats) in the genomes of *Dictyostelium giganteum*, *D. discoideum*, and *D. firmibasis*. The y-axis is plotted on a logarithmic scale to accommodate differences in repeat abundance across motif classes. Mononucleotide repeats are the most abundant class in all three genomes, followed by tri- and dinucleotide repeats, whereas longer motifs (tetra- to hexanucleotide repeats) occur at substantially lower frequencies. Numbers above bars indicate absolute SSR counts for each motif class and species.

### Simple sequence repeat (SSR) profiling in *Dictyostelium giganteum*

In addition to transposable elements, microsatellite composition was assessed. MISA-based SSR analysis revealed an AT-rich, microsatellite-dense genome in *D. giganteum*, with mononucleotide A/T tracts representing the predominant repeat class, followed by TA dinucleotides and AAT trinucleotides as the most abundant higher-order motifs. Tetra- and pentanucleotide SSRs were comparatively rare and mainly comprised AAAT and AAAAT motifs, respectively, whereas hexanucleotide repeats were dominated by AAAAAT with additional low-frequency variants, indicating a largely conserved but subtly polymorphic SSR landscape suitable for strain-level genotyping.

### Transcriptome analysis

To validate gene models and assess transcriptional activity, transcriptome data were integrated with the genome assembly. Transcriptome data were obtained from a *D. giganteum* isolate (NCBI BioProject Accession PRJNA48443; JGI GOLD ID Gp0008207) to validate our genomic assembly and annotation. De novo Trinity assembly of these sequences produced 18,542 transcript models, spanning 9.78 Mb with an average length of 527 bp and a maximum length of 7.8 kb. Mapping these transcripts to our 38.52 Mb reference genome revealed coverage of 25.38% of the assembly, indicating that approximately one-quarter of the genome is transcriptionally active under the sampled conditions. This analysis provided expression support for 5,549 annotated genes.

### Developmental Stage Assignment

Beyond structural validation, the transcriptome provided insight into the developmental context. Using Swiss-Prot functional annotation, we compared expressed transcripts with stage-specific markers defined for *Dictyostelium discoideum* (Loomis and Shaulsky 2011; Banu et al. 2026), surveying seven developmental stages from vegetative growth to culmination.

**Figure 4:**
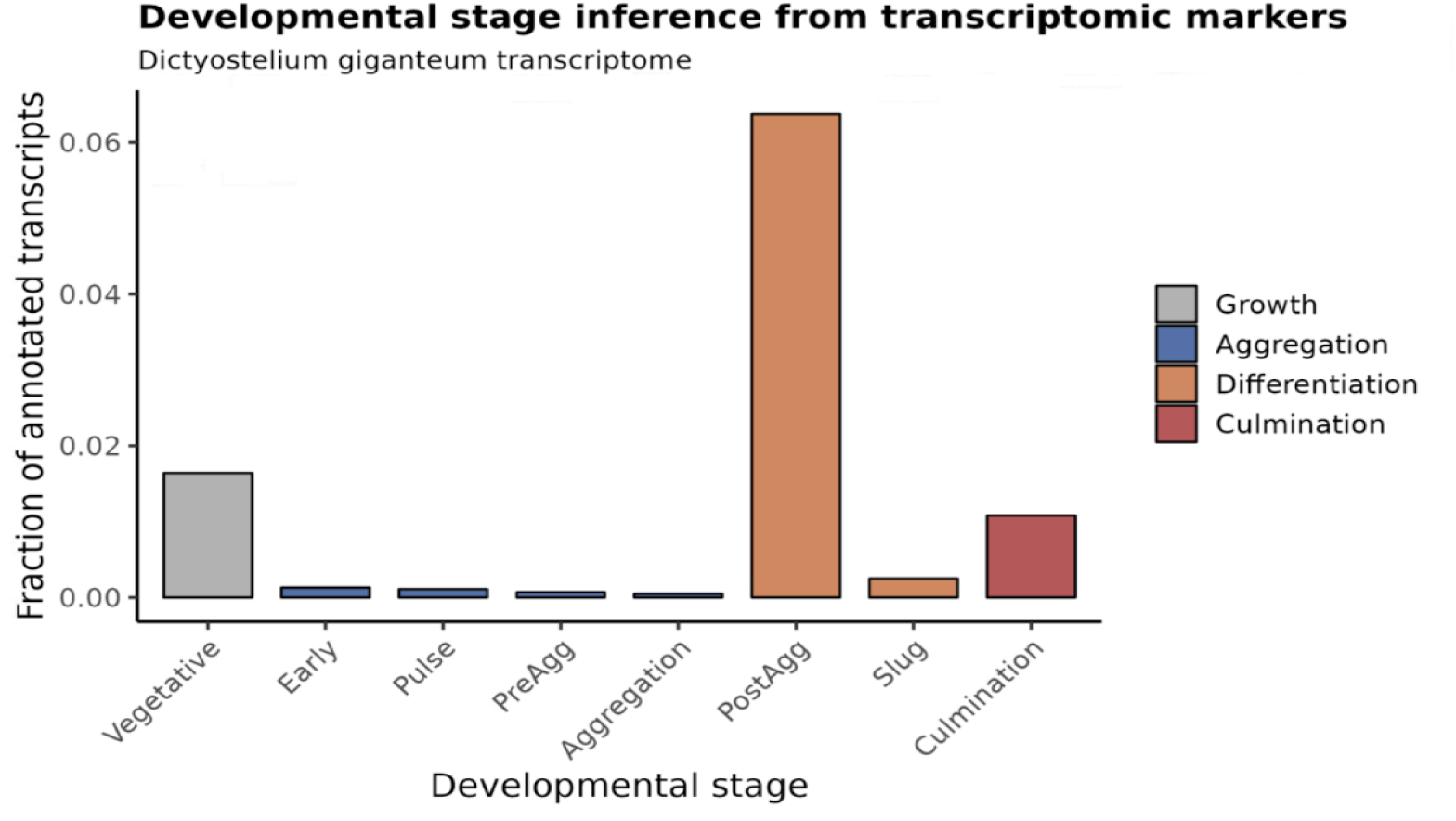
Developmental stage assignment using canonical transcriptomic markers: Distribution of stage-specific transcriptional markers across defined developmental phases of *Dictyostelium giganteum*, including vegetative growth, early starvation response, pulse response, aggregation, post-aggregation differentiation, slug formation, and culmination. Fractions represent normalized counts of Swiss-Prot annotated transcripts matching canonical marker genes for each stage. The dominance of post-aggregation markers, along with enrichment of culmination-associated genes and minimal representation of early developmental markers, supports assignment of the transcriptome to a late developmental stage encompassing slug formation and early culmination.

Markers of vegetative growth and early post-starvation phases were largely absent, excluding growth-phase or early starvation origins. In contrast, post-aggregation markers were strongly enriched, particularly those associated with slug formation and early culmination. Together, these findings indicate that the transcriptome predominantly represents late developmental stages, spanning slug formation and early culmination.

### Ribosomal DNA

Following transcriptomic validation, ribosomal DNA organization was examined. Ribosomal DNA (rDNA) in *Dictyostelium giganteum* is predominantly localized to unplaced, repeat-rich contigs and is absent from the chromosome-sized nuclear scaffolds, indicating that rRNA genes are not integrated into chromosomal DNA. Sequence comparison with the palindromic extrachromosomal rDNA element described in *D. discoideum* revealed extensive regions of high sequence identity, supporting a conserved extrachromosomal mode of rDNA organization. In total, the *D. giganteum* genome contains 16 rRNA genes, comparable to *D. discoideum* 15-18 rRNA genes.

**Figure 5A:**
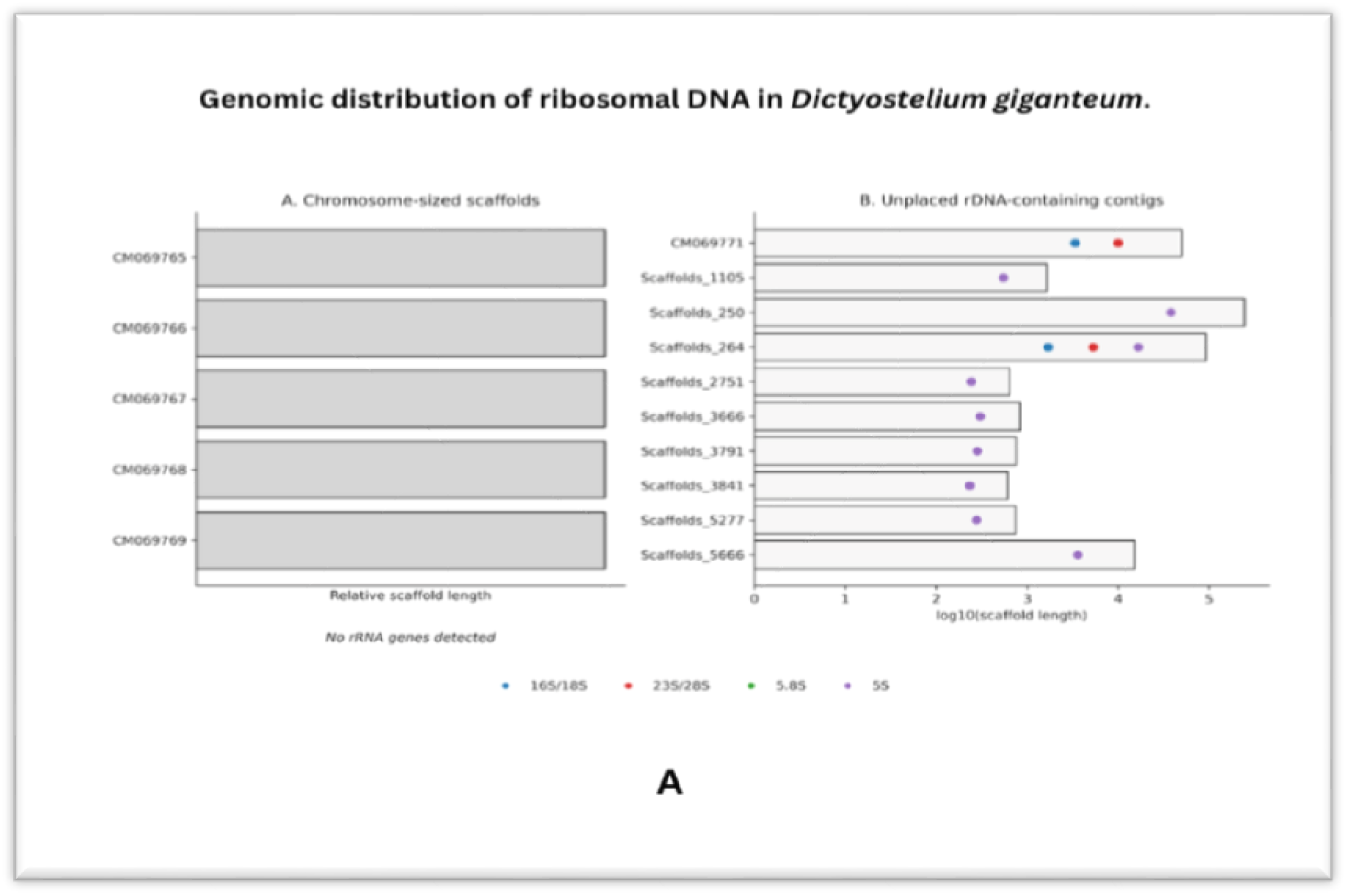
Genomic distribution of ribosomal DNA in *Dictyostelium giganteum*. Schematic representation of the genomic localization of ribosomal RNA (rRNA) genes in the *Dictyostelium giganteum* genome. **(A)** The five chromosome-sized nuclear scaffolds (CM069765–CM069769) are shown as long horizontal bars and contain no detectable rRNA gene annotations, indicating the absence of chromosomal rDNA arrays. **(B)** Unplaced scaffolds are shown as shorter bars of varying length (displayed on a logarithmic scale for visualization) and contain multiple rRNA gene annotations, including 16S/18S, 23S/28S, 5.8S, and 5S rRNAs, indicated by colored markers. rRNA genes are enriched on these repeat-rich, unplaced contigs and are absent from chromosome-sized scaffolds. This distribution is consistent with the maintenance of rDNA as extrachromosomal elements and with the fragmented assembly of palindromic rDNA structures, rather than integration into chromosomal DNA.

### tRNA genes

In addition to rRNA genes, the nuclear genome encodes an extensive tRNA repertoire. The *Dictyostelium giganteum* genome encodes 473 transfer RNA (tRNA) genes, representing all 20 standard amino acids as well as a single selenocysteine tRNA (tRNA-Sec) (Shrimali et al. 2005; Chan et al. 2021). Strand distribution is approximately balanced, with 262 genes encoded on the forward strand and 211 on the reverse strand.

To evaluate regulatory conservation, upstream promoter features were analyzed. B-box motifs were detected in 70.9% of upstream regions, while A/U-rich and TATA-like elements (Fig 5B) were present in the vast majority of loci, supporting assembly fidelity and functional conservation (Hofmann et al. 1991).

**Figure 5B:**
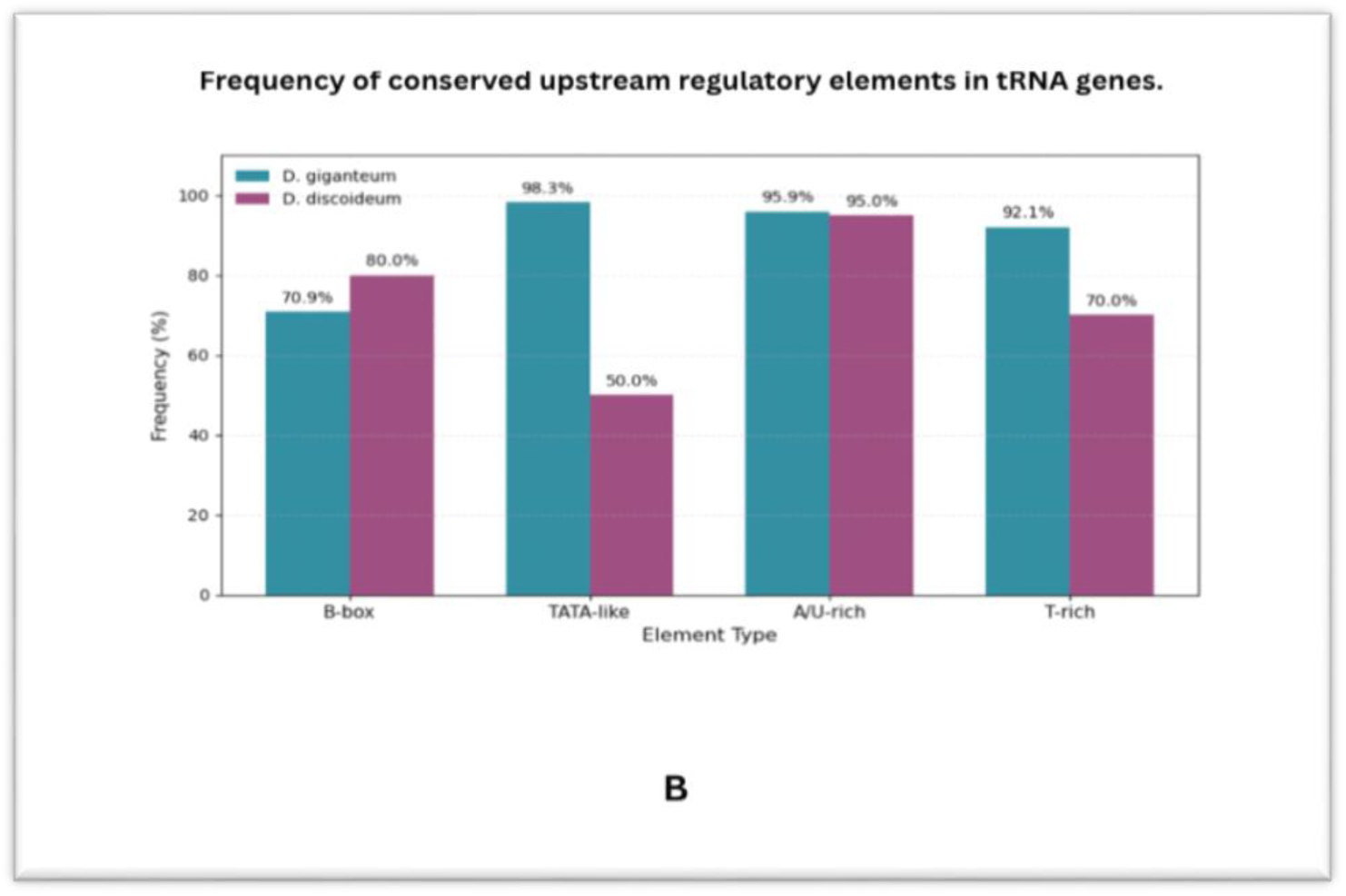
Frequency of conserved upstream regulatory elements in tRNA genes. Bar plot showing the proportion of tRNA genes containing B-box, TATA-like, A/U-rich, and T-rich upstream elements in *Dictyostelium giganteum* and *D. discoideum*. Percentages indicate the fraction of tRNA loci associated with each regulatory feature.

### Spliceosomal and all Nucleolar RNAs

Beyond tRNA genes, additional non-coding RNAs were identified. Homology-based annotation using *D. discoideum* sequences identified a presumably complete set of major spliceosomal small nuclear RNAs (U1, U2, U4, U5, and U6) in *Dictyostelium giganteum*. These RNAs show high sequence conservation and preserved predicted secondary structures relative to their counterparts in *D. discoideum*, indicating strong evolutionary conservation of the core spliceosomal machinery (Aspegren 2004; Hinas et al. 2006).

In addition, a diverse repertoire of small nucleolar RNAs (snoRNAs) was identified, predominantly belonging to the DdR family, including DdR2, DdR4, DdR5, DdR6, DdR10, DdR15, DdR16, and DdR18. Many of these snoRNAs display clear one-to-one correspondence with orthologous loci, consistent with conserved roles in ribosomal RNA modification and processing.

### DUSE-Associated and Structured ncRNAs

Complementing homology-based annotation, motif-based searches were performed. A motif-based genomic scan identified two loci containing the Dictyostelium upstream sequence element (DUSE; [AT]CCCA[AT]AA), a promoter motif previously characterized in *Dictyostelium discoideum*, located approximately 60 bp upstream of candidate non-coding RNA loci in *D. giganteum*. The associated downstream transcripts are predicted to form stable hairpin secondary structures, with minimum free energies of –9.2 and –7.6 kcal mol⁻¹, consistent with small structured RNAs. The presence of the DUSE promoter element, originally described in *D. discoideum*, together with stable RNA secondary structure, suggests that *D. giganteum* also transcribes promoter-linked structured ncRNAs analogous to those reported in *D. discoideum* (Hinas et al. 2006).

### Proteome Content and Organization

Following genome annotation, proteome composition was analyzed. The predicted proteome of *Dictyostelium giganteum* comprises 13,251 protein-coding genes encoding a total of 6.99 million amino acids. Predicted protein lengths range from fewer than 50 amino acids to several thousand amino acids. The mean protein length is 528.9 amino acids, comparable to that observed in other *Dictyostelium* species.

Protein length distribution analysis showed that approximately one-quarter of the *D. giganteum* proteome consists of proteins between 500 and 1,000 amino acids in length. Proteome size and length distributions are broadly similar to those reported for related Dictyostelium species.

### Genome Compactness and Codon Bias

Consistent with genome structure, coding density and nucleotide composition were examined. The *D. giganteum* genome is compact, with approximately 65.7% of the assembly consisting of protein-coding sequences. The genome exhibits pronounced AT-richness (76.3%), which influences codon usage patterns, with a preference for NNA and NNT codons.

Amino acids encoded by AT-rich codons—particularly asparagine, lysine, phenylalanine, and tyrosine—are enriched in the proteome, whereas amino acids encoded by GC-rich codons, including proline, arginine, alanine, and glycine, are proportionally reduced.

### Amino Acid Composition and Low-Complexity Features

In line with codon bias, the proteome exhibits elevated asparagine and glutamine content. Consistent with the AT-driven codon composition, the *D. giganteum* proteome contains a high proportion of asparagine (N) and glutamine (Q) residues. However, the organization of these residues differs from *D. discoideum* (Table 2). Long poly-N/poly-Q tracts (≥20 amino acids) were identified in 6.5% of *D. giganteum* proteins, compared with approximately 17% in *D. discoideum (Malinovska et al., 2015)*.

**Table 2:** **Low-Complexity and Repeat Composition in *D. giganteum***

| Feature | Count | % of Proteome |
| --- | --- | --- |
| Proteins with homopolymers $\geq 10$ aa | 3,472 | 21.8% |
| Proteins with polyN/polyQ $\geq 20$ aa | 1,034 | 6.5% |
| • Poly-asparagine (N $\geq 20$ aa) | 714 | — |
| • Poly-glutamine (Q $\geq 20$ aa) | 320 | — |
| Proteins with dipeptide repeats $\geq 15$ aa | 3,283 | 20.6% |
| Proteins with any repeat (homo- or dipeptide) | 4,003 | 25.1% |

Low-complexity sequences are common in the *D. giganteum* proteome, but long homopolymeric N/Q expansions occur less frequently than in *D. discoideum*.

### Functional Annotation and Domain Organization

To further characterize proteome function, domain-level annotation was performed. Functional annotation using eggNOG-mapper assigned COG categories, Gene Ontology terms, KEGG pathways, and Pfam domains to 13,251 predicted proteins (>98% coverage). COG analysis showed that 44.2% of proteins were classified as poorly characterized (COG S). Among characterized categories, post-translational modification machinery (COG O; 6.5%) and signal transduction components (COG T; 5.4%) were the most represented functional groups.

### Pfam domain architecture emphasizes signalling and metabolic versatility

Pfam annotation identified 3,790 distinct domain types across 19,728 total occurrences (Supplementary Figure 1A). FNIP repeats were the most abundant domain class (419 proteins; 22.6%), followed by protein kinase domains (258 proteins; 13.9%). Additional prominent domain families included small GTPases (Ras family), WD repeats, ankyrin repeats, IPT/TIG domains, ABC transporters, cytochrome P450 domains, RNA recognition motifs (RRM), and helicases. Collagen triple-helix repeats (PF01391) were detected in *D. discoideum* and *D. giganteum* but were not identified in *D. firmibasis*, occurring at low copy numbers. Several Dictyostelium-specific domain families—including the Dicty repeat (PF00526), counting factor domain (PF11912), and HssA/B-related domain (PF05710)—were present across Dictyostelium species but were not detected in representative metazoan genomes. Notably, FNIP-domain–containing proteins have also been reported in *Entamoeba histolytica*, where comparative genomic analysis identified FNIP repeats as part of the limited set of Amoebozoa-specific gene families shared between *Dictyostelium* and *Entamoeba* (Song et al. 2005).

### Gene family distribution: -

Orthologous clustering identified 5,483 gene families encompassing 71.9% of the proteome (Supplementary Figure 1B). Protein kinases (∼400 members) constitute the largest gene family. Polyketide synthases are also represented by multiple family members.

Gene families containing RhoGAPs, RhoGEFs, and Rho GTPases were identified, along with expanded families of heat shock proteins, ABC transporters, and myosins. Overall gene family size distributions show broad similarity to those observed in *D. discoideum* (Supplementary Figure 1B).

### Orthogroup Architecture Reveals Extensive Dictyostelium-Specific Innovation

To assess evolutionary conservation, comparative orthology analysis was conducted. Comparative orthology analysis across *Dictyostelium discoideum*, *D. giganteum*, *D. firmibasis*, and *Entamoeba* resolved a hierarchical pattern of gene conservation within Amoebozoa (Figure 6). A total of 1,612 orthogroups were conserved across all four species, defining a deeply shared Amoebozoan core enriched for fundamental cellular functions.

**Figure 6.**
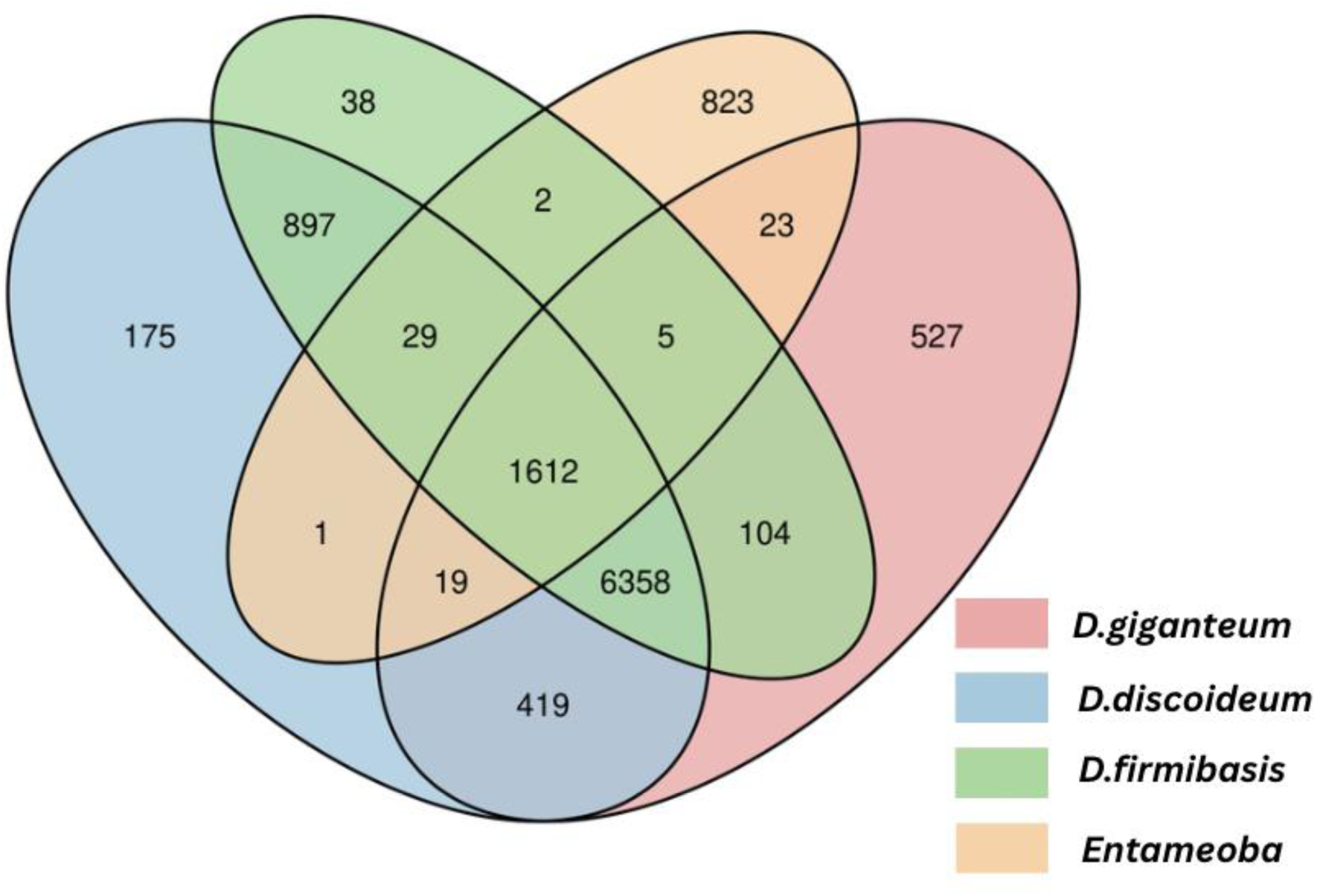
Orthogroup sharing among three Dictyostelium species and *Entamoeba*. Orthogroup comparison across three Dictyostelium species and *Entamoeba* identified a large Dictyostelium-specific core comprising 6,358 orthogroups absent from *Entamoeba*, alongside 1,612 deeply conserved orthogroups shared across all four Amoebozoa. Species-restricted repertoires were most pronounced in *Entamoeba* (823 orthogroups) and *D. giganteum* (527), suggesting lineage-specific innovation and potential adaptation to parasitism or ecological specialization. Extensive pairwise sharing among Dicty species, particularly between *D. discoideum* and *D. firmibasis* (897 orthogroups), highlights strong genus-level conservation. Orthogroups shared between *Entamoeba* and pairs of Dicty species but absent from the third Dicty species were comparatively few (5–29 orthogroups per combination), consistent with substantial divergence between parasitic and social amoebozoan lineages.

In contrast, 6,358 orthogroups were shared exclusively among the three *Dictyostelium* species but were absent from *Entamoeba*, representing a substantial genus-specific gene repertoire associated with social amoebae.

Lineage-restricted innovation was also evident, with 175 orthogroups unique to *D. discoideum*, 38 to *D. firmibasis*, 527 to *D. giganteum*, and 823 unique to *Entamoeba*, reflecting ongoing species-level diversification following divergence.

### Evidence that the “Metazoan Toolkit” Predates Metazoa

Building on orthology results, domain-level comparisons with metazoans were performed. Previous comparative studies have suggested that several components of the developmental and signalling machinery of animals predate Metazoa (Brunet and King 2017; Elizabeth Pennisi 2018; Nanjundiah et al. 2018). Using *Entamoeba* and Dictyostelium species (*D. discoideum*, *D. firmibasis*, and *D. giganteum*) as non-metazoan references, we performed a systematic domain-level comparison with representative metazoans to enable lineage-wide assessment within Dictyostelia (Figure 7).

**Figure 7.**
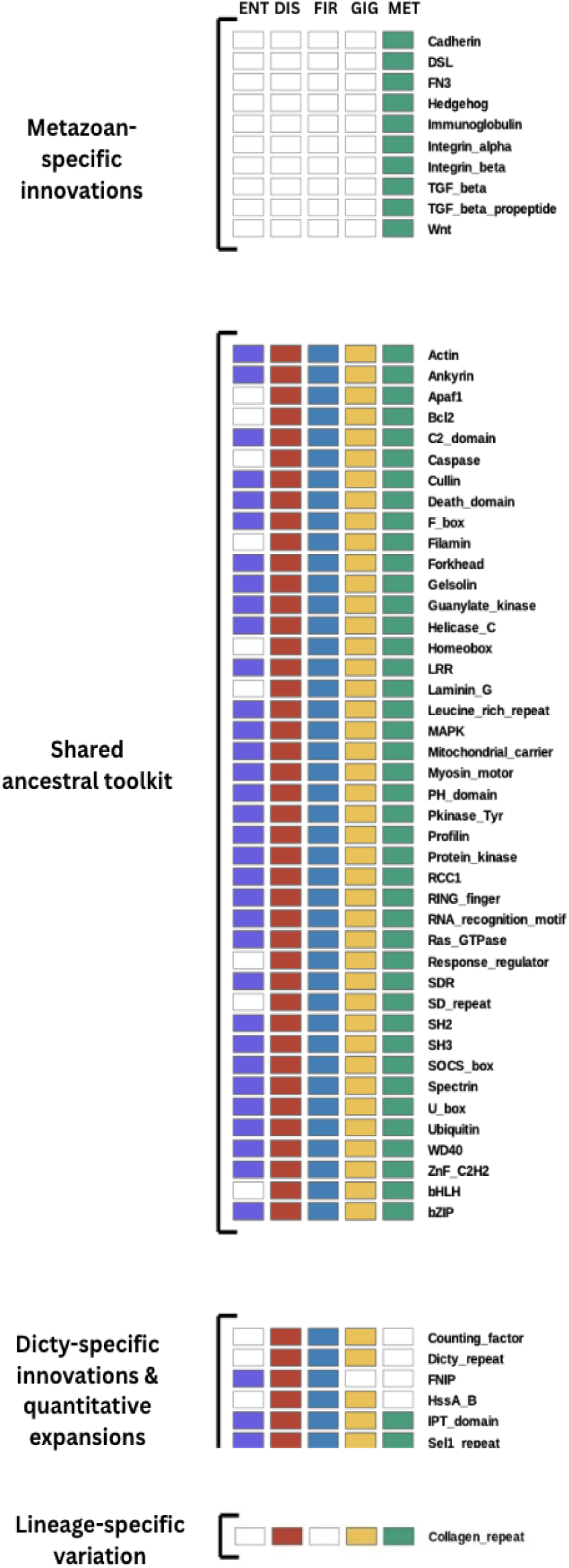
Evolutionary distribution of multicellularity-associated protein domains across non-social amoebozoans, social Dictyostelia, and Metazoa. Presence–absence matrix of selected Pfam domains in the non-social amoebozoan *Entamoeba* (ENT), the social amoebozoans *Dictyostelium discoideum* (DIS), *D. firmibasis* (FIR), *D. giganteum* (GIG), and representative Metazoa (MET). Filled boxes indicate presence; empty boxes indicate absence. Domains are grouped by evolutionary distribution. Metazoan-specific innovations (e.g., Cadherin, Immunoglobulin, Integrins, Wnt, Hedgehog, TGF-β) are restricted to animals. The shared ancestral toolkit comprises conserved intracellular signalling, cytoskeletal, transcriptional, and ubiquitin-regulatory modules (e.g., Protein kinase, Ras, SH2/SH3, WD40, Actin, MAPK) present in both amoebozoans and metazoans, indicating deep eukaryotic origins. Dictyostelium-specific innovations and quantitative expansions include lineage-restricted repeats and amplified ancestral domains associated with social development. Lineage-specific variation highlights heterogeneous retention among Dictyostelium species. Together, these patterns support independent origins of aggregative and clonal multicellularity from a shared pre-metazoan regulatory repertoire.

Most protein domains commonly associated with metazoan development and signalling are present in *D. giganteum* in substantial copy numbers (Figure 7). These include kinase domains (tyrosine kinases and serine/threonine kinases), signalling modules such as SH2/SH3 domains and Ras/Rho GTPases, scaffolding proteins including WD40 repeats and leucine-rich repeats, major transcription factor families (homeobox, forkhead, Myb, bHLH), apoptosis-related domains such as caspases, and core cytoskeletal components (actin, myosin, spectrin, gelsolin). The presence of these domain classes is consistent with earlier analyses of unicellular relatives of animals (Richter and King 2013; Brunet and King 2017). Domain-level reconstructions have also reported that many of these modules were established prior to the emergence of animals (Xiao et al. 2025). In several cases, these domain families are represented by multiple copies in *D. giganteum*.

In contrast, several canonical metazoan extracellular and structural domains were not detected in *D. giganteum* (Figure 7). These include classical cadherin domains, integrin α and β extracellular domains, TGF-β ligand domains, Hedgehog domains, Notch DSL domains, zona pellucida domains, and collagen triple-helix repeats. Comparative studies of early animal evolution have documented expansion of extracellular adhesion systems and morphogen-based signalling pathways within Metazoa (Newman and Bhat 2008; Brunet and King 2017; Xiao et al. 2025). Although *D. giganteum* lacks a classical cadherin homolog, 61 proteins containing catenin domains were identified, including homologs of both β- and α-catenin (Weis et al. 2013).

Notably, a subset of signalling and cytoskeletal modules present in Dictyostelium and Metazoa but reduced or absent in *Entamoeba* may represent components associated with the transition from solitary amoebae to aggregative multicellularity within Dictyostelia.

### Horizontal Gene Transfer Candidates

In addition to vertical inheritance, potential horizontal gene transfer events were investigated. To investigate the extent of horizontal gene transfer (HGT) in *Dictyostelium giganteum*, we examined Pfam domains and phyletic signatures indicative of bacterial origin. This analysis identified multiple domains consistent with horizontally acquired genes.

Several HGT-derived domains previously described in *D. discoideum* (Eichinger et al. 2005) are also present in *D. giganteum* (Supplementary Table S3), including ThyX, TerD, OsmC, polyphosphate kinase (PP_kinase), peptidase M15/S13, Endotoxin_N, DUF885, DUF1121, DUF1289, and DUF1294. These domains are associated with functions such as thymidylate synthesis, detoxification, oxidative stress response, polyphosphate metabolism, and bacterial cell-wall degradation.

A subset of HGT candidates reported in *D. discoideum*—including CnaB, Dyp_peroxidase, and luc4/LucC—were not detected in *D. giganteum*. Several candidate genes identified in *D. giganteum* show the closest sequence similarity to members of the Burkholderiaceae (DiSalvo et al. 2015).

### Orthologs of genes associated with Human Disease

To assess the capacity of *Dictyostelium giganteum* to serve as a model system for studying human disease genes, we performed a systematic orthology search using the predicted *D. giganteum* proteome. A total of 9,900 confirmed human disease-associated protein sequences were used as queries in BLASTP searches against the *D. giganteum* protein set. Using a stringent e-value cutoff of 1e−40, we identified 247 *D. giganteum* proteins with strong homology to human disease genes.

These orthologues span multiple disease-relevant categories, including DNA repair (e.g., MSH2, MLH1, PMS2), neurological and lysosomal disorders (e.g., LIS1, HEXA), and metabolic deficiencies (e.g., G6PD). For many orthologs, similarity extended across the majority of the protein length, with up to ∼90% identity for conserved cytoskeletal and enzymatic proteins.

Comparison with a curated panel of 30 validated *D. discoideum* disease-gene orthologs (Pears et al. 2021) showed that 19 (63.3%)(Table 3) are also present in *D. giganteum*, whereas several of these genes are absent from both *Saccharomyces cerevisiae* and *Schizosaccharomyces pombe*.

**Table 3:** **Human disease–related genes conserved in *Dictyostelium giganteum***

| Gene | Disease Category | Human Ortholog / Accession |
| --- | --- | --- |
| **MSH2** | Cancer | sp\|Q553L4\|MSH2_DICDI |
| **MLH1** | Cancer | sp\|Q54KD8\|MLH1_DICDI |
| **PMS2** | Cancer | sp\|P54279\|PMS2_MOUSE |
| **XPE** | Xeroderma pigmentosum | sp\|B0XPE7\|BTAF1_ASPFC |
| **XPF** | Xeroderma pigmentosum | sp\|Q54PN5\|XPF_DICDI |
| **RASH** | Oncogene (RAS) | sp\|P23175\|RASH_MSVNS |
| **CDK4** | Cell cycle regulation | sp\|Q91727\|CDK4_XENLA |
| **OCRL** | Neurological disorder | sp\|D3ZGS3\|OCRL_RAT |
| **LIS1** | Neurological disorder | sp\|D3BUN1\|LIS1_HETP5 |
| **ALD** | Neurological disorder | sp\|Q54UP4\|TALDO_DICDI |
| **HEXA** | Neurological disorder | sp\|P13723\|HEXA1_DICDI |
| **PPT1** | Neurological disorder | sp\|Q8HXW6\|PPT1_MACFA |
| **NPC1** | Niemann–Pick disease | sp\|O35604\|NPC1_MOUSE |
| **CBPD** | Periodontitis | sp\|P83852\|CBPD_LOPSP |
| **G6PD** | Haematological / immune disorder | sp\|Q557D2\|G6PD_DICDI |
| **C24B** | Chronic granulomatous disease | sp\|O95487\|SC24B_HUMAN |
| **DTD** | Metabolic disorder | sp\|Q54EY1\|DTD_DICDI |
| **DRA** | Metabolic disorder | sp\|Q54Y26\|NDRA_DICDI |
| **ATP2** | Metabolic disorder | sp\|Q55CA5\|ATP23_DICDI |

## V. DISCUSSION

### Overview and Assembly Strategy

*Dictyostelium* occupies a unique evolutionary position as a social amoeba bridging unicellular and multicellular life, making it a powerful model for studying the origins of complex cellular communication, differentiation, and developmental regulation. To provide a genomic foundation for these investigations, we generated a species-level consensus genome of *Dictyostelium giganteum* using an integrated strategy combining de novo assembly, reference-guided scaffolding, iterative polishing, and progressive multi-strain alignment. Independent assemblies from six wild strains were aligned against a high-quality backbone, enabling incorporation of conserved high-identity regions while minimizing strain-specific artifacts and preserving chromosomal structure.

The final assembly spans **38.52 Mb**, with an **N50 of 3.01 Mb**, **L50 of 5**, and ∼**343× coverage**, resolving into five chromosome-scale scaffolds consistent with previous cytogenetic observations (Kumar et al. 2024). The high completeness (BUSCO >92% complete genes) supports its reliability as a genomic framework.

Although *D. discoideum* and *D. firmibasis* possess six chromosomes (Eichinger et al. 2005; Edelbroek et al. 2024), *D. giganteum* has only five chromosomes. Whole-genome synteny demonstrates that genomic content corresponding to the sixth chromosome is not absent but redistributed across the five scaffolds. Therefore, the evolution of chromosomes in Dictyostelia is mediated by large-scale genomic rearrangements or fusions, rather than gene loss. DNA sequence similarity at the DNA level also confirms the relationships: sequence identity varies from ∼22% for *D. discoideum* to ∼40% for *D. firmibasis*, quantitatively validating the evolutionary proximity of *D. giganteum* and *D. firmibasis* (Schaap et al. 2006).

### Genome Architecture and Base Composition

The genome is highly compact, with **65.7% protein-coding sequence**, and exhibits extreme **AT richness (∼76%)**, a defining feature of dictyostelid genomes. CpG dinucleotides are significantly depleted (observed/expected ratio **0.565**), indicating moderate-to-strong CpG suppression. However, the lack of canonical DNA methyltransferases suggests that this depletion is likely a consequence of long-term mutational bias associated with AT enrichment rather than active genomic methylation.

Repetitive sequences comprise approximately **18%** of the nuclear genome (∼5.2 Mb). Among transposable elements, DIRS1 elements are the most conserved and abundant, while most other TE families are present as fragmented copies, indicating historical rather than ongoing expansion. Simple sequence repeat analysis reveals a microsatellite-dense genome dominated by A/T mononucleotide tracts, consistent with overall base composition. Such polymorphic SSRs may provide useful markers for strain-level genotyping in natural populations.

### Non-Coding RNAs and Ribosomal DNA Organization

The *D. giganteum* genome encodes a large complement of non-coding RNAs, including tRNAs, spliceosomal RNAs, snoRNAs, and candidate structured ncRNAs associated with the DUSE promoter element. Identification of a full complement of major spliceosomal snRNAs (U1–U6) and a conserved set of DdR-family snoRNAs supports broad conservation of core RNA processing machinery across Dictyostelia.

Sixteen rRNA genes were identified, similar to the 15–18 copies reported in *D. discoideum* supporting conservation of overall rDNA dosage. Ribosomal DNA is absent from chromosome-scale scaffolds and instead localized to repeat-rich, unplaced contigs, explained by partial assembly of a palindromic extrachromosomal structure, which is difficult to resolve with short reads. This distribution is consistent with extrachromosomal, palindromic rDNA elements rather than chromosomally integrated arrays. Such organization may allow flexible regulation of rRNA gene dosage and suggests that chromosomal nucleolar organizer regions seen in many eukaryotes could represent a derived state.

The genome encodes 473 tRNA genes, including a functional selenocysteine tRNA, and displays strong conservation of upstream regulatory motifs—B-box elements in approximately 71% of loci and TATA-like elements in over 95%—closely mirroring the pattern observed in *D. discoideum*.

### Protein-Coding Gene Complement and Annotation Support

The proteome prediction includes 13,251 protein-coding genes with an average protein size of about 529 amino acids. AT-driven codon bias predictably leads to an enrichment of asparagine and glutamine residues, but extended polyN/polyQ tracts of 20 or more amino acids are found in only 6.5% of proteins—well below the approximately 17% found in *D. discoideum* (Malinovska et al. 2015). This does not indicate a general reduction of N/Q residues, but rather a more even distribution of these residues with fewer extreme homopolymeric tracts, which could decrease the propensity for aggregation while maintaining the regulatory plasticity of disordered regions or maybe. The predicted gene number in *D. giganteum* is comparable to that of *D. discoideum* and exceeds that of *D. firmibasis*. Functional annotation using domain and orthology-based methods assigns putative functions to the vast majority of predicted proteins (>98%). However, a substantial fraction remains poorly characterized, consistent with extensive lineage-specific innovation across dictyostelids.

Transcriptome mapping supports the assembly and annotation, providing expression evidence for **5,549 genes**. Developmental marker analysis indicates that the transcriptome predominantly represents late developmental stages (slug formation and early culmination), with enrichment of spore coat proteins and culmination-associated genes. This confirms conservation of the multicellular developmental program across dictyostelids.

### Comparative Context and Evolutionary Implications

Functional and orthogroup comparisons combined provide strong support for a consistent view: evolutionary innovation in Dictyostelium is a result of the expansion and diversification of gene families from a common ancestor rather than the emergence of entirely novel systems. A set of 1,612 orthogroups is conserved across Dictyostelium species and Entamoeba (parasitic); which served not only as a non-social control but also as an outgroup to bracket dictyostelid-enriched gene families. A large number (6,358) of orthogroups are specific to Dictyostelium, reflecting innovations tied to social behaviour. A further 527 orthogroups appear unique to *D. giganteum*, pointing to species-level specialisation.

Domain-level annotation highlights FNIP repeats and protein kinase domains as the most abundant Pfam families, consistent with the importance of nutrient sensing and phosphorylation-based signalling in developmental regulation. Collagen triple-helix repeats were detected in *D. giganteum* and *D. discoideum* but not *D. firmibasis*, while Dictyostelium-specific families—including the Dicty repeat, counting factor domain, and HssA/B-related domain—are present across all three Dictyostelium species but absent from Metazoa, pointing to lineage-specific innovations associated with aggregative multicellularity.

Many intracellular domains associated with the metazoan toolkit—protein kinases, Ras/Rho GTPases, SH2/SH3 domains, major transcription factor families, apoptosis regulators, and core cytoskeletal components—are present in *D. giganteum*, often in expanded copy numbers. In contrast, canonical extracellular adhesion and morphogen-signalling domains characteristic of animals—cadherins, integrins, TGF-β, Hedgehog, and Notch DSL—are absent. Notably, 61 catenin-domain proteins were identified despite the absence of classical cadherins, suggesting that the intracellular adhesion and polarity machinery predates the evolution of cadherin-based extracellular adhesion. Together, these observations support a model in which animal multicellularity arose through the elaboration of pre-existing intracellular regulatory systems combined with the later addition of extracellular innovations.

### Horizontal Gene Transfer Candidates and Microbiome Associations

Analysis of the *Dictyostelium giganteum* genome identified several proteins containing domains typically associated with bacteria, including ThyX, OsmC, polyphosphate kinase, and siderophore-related proteins. Sequence similarity searches indicated that many of these proteins show closest similarity to members of the *Burkholderiaceae*. Furthermore, amoeba-associated *Burkholderia* were observed to establish farming symbiosis with naïve amoeba hosts (DiSalvo et al. 2015).

Finally, comparison with human disease-associated genes identified **247 strong orthologues**, including conserved DNA repair and cytoskeletal proteins. Notably, 19 of 30 validated *D. discoideum* disease-gene orthologues (Pears et al. 2021) are also present in *D. giganteum*, whereas several are absent from yeast models. This highlights the unique evolutionary position of Dictyostelia as a bridge between unicellular fungi and multicellular animals.

Taken together, the *D. giganteum* genome reveals a lineage that is structurally dynamic yet functionally conserved, combining chromosomal plasticity, extreme compositional bias, and lineage-specific innovations with deep conservation of the intracellular regulatory machinery shared with animals. Rather than representing a structural intermediate between unicellular and multicellular life*, D. giganteum* illustrates how complex multicellular behaviour can arise from the development and coordination of existing regulatory networks. In this respect, it holds a key evolutionary position, bridging solitary eukaryotes and the complex multicellularity of animals, providing a glimpse into the ancestral regulatory toolkit from which animal complexity ultimately arose.

### Mitochondrial Genome Conservation and Divergence

The *D. giganteum* mitochondrial genome assembly measures **50,452 bp**, encodes **36 protein-coding genes**, and maintains highly polarized gene organization, with nearly all ORFs located on a single strand. Despite minor size differences among dictyostelids, mitochondrial architecture remains broadly conserved, suggesting stronger structural constraints at the organellar level. A notable feature is strong strand polarization, with nearly all open reading frames on a single strand. This may reflect lineage-specific constraints on replication or transcription, although its mechanistic implications remain unclear. The mitochondrial genome has a slightly higher protein-coding fraction (∼67.9%) than the nuclear genome, consistent with compact mitochondrial architecture.

## CONCLUSION

The consensus genome of *Dictyostelium giganteum* presented here provides a high-quality genomic framework for understanding the molecular basis of aggregative multicellularity within Dictyostelia. Despite AT-richness, chromosomal rearrangements, and lineage-specific innovations, the genome reveals deep conservation of intracellular regulatory and signalling machinery shared with other dictyostelids and, in part, with Metazoa. At the same time, the absence of canonical metazoan extracellular adhesion and morphogen-signalling domains reinforces the view that complex multicellular behaviour in Dictyostelium evolved through elaboration of pre-existing intracellular systems rather than through the adoption of animal-specific innovations. Comparative orthology highlights a substantial Dictyostelium-specific gene repertoire alongside a conserved Amoebozoan core, while evidence of bacterial-derived domains underscores ecological interactions that have shaped genome evolution. Together, these findings position *D. giganteum* as a key evolutionary reference within Amoebozoa, bridging solitary and social lifestyles and offering insight into how complex developmental programs can emerge from ancient eukaryotic regulatory toolkits.

## ACKNOWLEDGEMENTS & FUNDING

Illumina reads were generated using services from the BioIT centre at Institute of Bioinformatics and Applied Biotechnology. The authors thank the Government of Karnataka for funding for sequencing and data analysis personnel via a BioIT grant and computing infrastructure via the Department of Information Technology, Biotechnology and Science and Technology. The authors also acknowledge using computing resources obtained as part of DBT Builder Sanction no. BT/INF/22/SP45402/2022 dated March 8, 2022, and DST-FIST Sanction no. SR/FST/LSI-536/2012.

## REFERENCES

Alonge M, Lebeigle L, Kirsche M, Jenike K, Ou S, Aganezov S, Wang X, Lippman ZB, Schatz MC, Soyk S. 2022. Automated assembly scaffolding using RagTag elevates a new tomato system for high-throughput genome editing. Genome Biol. 23:258.

Altschul SF, Gish W, Miller W, Myers EW, Lipman DJ. 1990. Basic local alignment search tool. J. Mol. Biol. 215:403–410.

Arias Del Angel JA, Nanjundiah V, Benítez M, Newman SA. 2020. Interplay of mesoscale physics and agent-like behaviors in the parallel evolution of aggregative multicellularity. EvoDevo 11:21.

Aspegren A. 2004. Novel non-coding RNAs in Dictyostelium discoideum and their expression during development. Nucleic Acids Res. 32:4646–4656.

Banu S, Anusha PV, Beltran-Alvarez P, Idris MM, Wollenberg Valero KC, Rivero F. 2026. The Proteome of Dictyostelium discoideum Across Its Entire Life Cycle Reveals Sharp Transitions Between Developmental Stages. Proteomes 14:3.

Bao W, Kojima KK, Kohany O. 2015. Repbase Update, a database of repetitive elements in eukaryotic genomes. Mob. DNA 6:11.

Basu S, Fey P, Jimenez-Morales D, Dodson RJ, Chisholm RL. 2015. dicty B ase 2015: Expanding data and annotations in a new software environment. genesis 53:523–534.

Bolger AM, Lohse M, Usadel B. 2014. Trimmomatic: a flexible trimmer for Illumina sequence data. Bioinformatics 30:2114–2120.

Bonner JT. 2003. On the origin of differentiation. J. Biosci. 28:523–528.

Bonner JT. 2009. The social amoebae: the biology of cellular slime molds. Princeton: Princeton University Press

Brefeld O. 1884. Botanische Untersuchungen über Myxomyceten und Entomophthoreen: Polysphondylium violaceum und Dictyostelium mucoroides. Conidiobolus utriculosus und minor. Verlag von Arthur Felix

Brown MW, Silberman JD. 2013. The Non-dictyostelid Sorocarpic Amoebae. In: Romeralo M, Baldauf S, Escalante R, editors. Dictyostelids. Berlin, Heidelberg: Springer Berlin Heidelberg. p. 219–242. Available from: http://link.springer.com/10.1007/978-3-642-38487-5_12

Brunet T, King N. 2017. The Origin of Animal Multicellularity and Cell Differentiation. Dev. Cell 43:124–140.

Camacho C, Coulouris G, Avagyan V, Ma N, Papadopoulos J, Bealer K, Madden TL. 2009. BLAST+: architecture and applications. BMC Bioinformatics 10:421.

Chan PP, Lin BY, Mak AJ, Lowe TM. 2021. tRNAscan-SE 2.0: improved detection and functional classification of transfer RNA genes. Nucleic Acids Res. 49:9077–9096.

DiSalvo S, Haselkorn TS, Bashir U, Jimenez D, Brock DA, Queller DC, Strassmann JE. 2015. *Burkholderia* bacteria infectiously induce the proto-farming symbiosis of *Dictyostelium* amoebae and food bacteria. Proc. Natl. Acad. Sci. [Internet] 112. Available from: https://pnas.org/doi/full/10.1073/pnas.1511878112

Edelbroek B, Kjellin J, Jerlström-Hultqvist J, Koskiniemi S, Söderbom F. 2024. Chromosome-level genome assembly and annotation of the social amoeba Dictyostelium firmibasis. Sci. Data 11:678.

Eichinger L, Pachebat JA, Glöckner G, Rajandream M-A, Sucgang R, Berriman M, Song J, Olsen R, Szafranski K, Xu Q, et al. 2005. The genome of the social amoeba Dictyostelium discoideum. Nature 435:43–57.

Elizabeth Pennisi EP. 2018. The momentous transition to multicellular life may not have been so hard after all. Sci. AAAS [Internet]. Available from: https://www.science.org/content/article/momentous-transition-multicellular-life-may-not-have-been-so-hard-after-all

Escalante R, Cardenal-Muñoz E. 2019. The Dictyostelium discoideum model system. Int. J. Dev. Biol. 63:317–320.

Fey P, Gaudet P, Curk T, Zupan B, Just EM, Basu S, Merchant SN, Bushmanova YA, Shaulsky G, Kibbe WA, et al. 2009. dictyBase—a Dictyostelium bioinformatics resource update. Nucleic Acids Res. 37:D515–D519.

Grabherr MG, Haas BJ, Yassour M, Levin JZ, Thompson DA, Amit I, Adiconis X, Fan L, Raychowdhury R, Zeng Q, et al. 2011. Full-length transcriptome assembly from RNA-Seq data without a reference genome. Nat. Biotechnol. 29:644–652.

Gurevich A, Saveliev V, Vyahhi N, Tesler G. 2013. QUAST: quality assessment tool for genome assemblies. Bioinformatics 29:1072–1075.

Hinas A, Larsson P, Avesson L, Kirsebom LA, Virtanen A, Söderbom F. 2006. Identification of the Major Spliceosomal RNAs in *Dictyostelium discoideum* Reveals Developmentally Regulated U2 Variants and Polyadenylated snRNAs. Eukaryot. Cell 5:924–934.

Hofmann J, Schumann G, Borschet G, Gösseringer R, Bach M, Bertling WM, Marschalek R, Dingermann T. 1991. Transfer RNA genes from Dictyostelium discoideum are frequently associated with repetitive elements and contain consensus boxes in their 5′ and 3′-flanking regions. J. Mol. Biol. 222:537–552.

Hu W-S, Jiang L-L, Liu P, Zhang X-Y, Wei W, Du X-H. 2024. Morphological and Phylogenetic Analyses Reveal Dictyostelids (Cellular Slime Molds) Colonizing the Ascocarp of Morchella. J. Fungi 10:678.

Huerta-Cepas J, Forslund K, Coelho LP, Szklarczyk D, Jensen LJ, Von Mering C, Bork P. 2017. Fast Genome-Wide Functional Annotation through Orthology Assignment by eggNOG-Mapper. Mol. Biol. Evol. 34:2115–2122.

Jones P, Binns D, Chang H-Y, Fraser M, Li W, McAnulla C, McWilliam H, Maslen J, Mitchell A, Nuka G, et al. 2014. InterProScan 5: genome-scale protein function classification. Bioinformatics 30:1236–1240.

Kalvari I, Nawrocki EP, Ontiveros-Palacios N, Argasinska J, Lamkiewicz K, Marz M, Griffiths-Jones S, Toffano-Nioche C, Gautheret D, Weinberg Z, et al. 2021. Rfam 14: expanded coverage of metagenomic, viral and microRNA families. Nucleic Acids Res. 49:D192–D200.

Kaushik S, Katoch B, Nanjundiah V. 2006. Social behaviour in genetically heterogeneous groups of Dictyostelium giganteum. Behav. Ecol. Sociobiol. 59:521–530.

Kumar R, S. Kulshreshtha P, Nanjundiah V, S. Kadandale J. 2024. The Chromosomes of Dictyostelium Giganteum. J. Chromosom. [Internet] 1. Available from: https://openaccesspub.org/chromosomes/article/2143

Li D, Liu C-M, Luo R, Sadakane K, Lam T-W. 2015. MEGAHIT: an ultra-fast single-node solution for large and complex metagenomics assembly via succinct de Bruijn graph. Bioinformatics 31:1674–1676.

Li H, Handsaker B, Wysoker A, Fennell T, Ruan J, Homer N, Marth G, Abecasis G, Durbin R, 1000 Genome Project Data Processing Subgroup. 2009. The Sequence Alignment/Map format and SAMtools. Bioinformatics 25:2078–2079.

Loomis WF, Shaulsky G. 2011. Developmental changes in transcriptional profiles. Dev. Growth Differ. 53:567–575.

Malinovska L, Palm S, Gibson K, Verbavatz J-M, Alberti S. 2015. Dictyostelium discoideum has a highly Q/N-rich proteome and shows an unusual resilience to protein aggregation. Proc. Natl. Acad. Sci. U. S. A. 112:E2620–2629.

Manni M, Berkeley MR, Seppey M, Simão FA, Zdobnov EM. 2021. BUSCO update: novel and streamlined workflows along with broader and deeper phylogenetic coverage for scoring of eukaryotic, prokaryotic, and viral genomes. Mol. Biol. Evol. 38:4647–4654.

Marçais G, Delcher AL, Phillippy AM, Coston R, Salzberg SL, Zimin A. 2018. MUMmer4: A fast and versatile genome alignment system. Darling AE, editor. PLOS Comput. Biol. 14:e1005944.

Nanjundiah V, Ruiz-Trillo I, Kirk D. 2018. Protists and multiple routes to the evolution of multicellularity. In: Cells in evolutionary biology. CRC Press. p. 71–118.

Nawrocki EP, Eddy SR. 2013. Infernal 1.1: 100-fold faster RNA homology searches. Bioinformatics 29:2933–2935.

NCBI Resource Coordinators, Agarwala R, Barrett T, Beck J, Benson DA, Bollin C, Bolton E, Bourexis D, Brister JR, Bryant SH, et al. 2018. Database resources of the National Center for Biotechnology Information. Nucleic Acids Res. 46:D8–D13.

Newman SA, Bhat R. 2008. Dynamical patterning modules: physico-genetic determinants of morphological development and evolution. Phys. Biol. 5:015008.

Niklas KJ, Newman SA eds. 2016. Multicellularity: Origins and Evolution. The MIT Press Available from: https://direct.mit.edu/books/book/4457/MulticellularityOrigins-and-Evolution

Pears CJ, Brustel J, Lakin ND. 2021. Dictyostelium discoideum as a Model to Assess Genome Stability Through DNA Repair. Front. Cell Dev. Biol. 9:752175.

Raper KB. 2014. The dictyostelids. Princeton University Press

Reddy AK, Balne PK, Garg P, Sangwan VS, Das M, Krishna PV, Bagga B, Vemuganti GK. 2010. Dictyostelium polycephalum infection of human cornea. Emerg. Infect. Dis. 16:1644.

Richter DJ, King N. 2013. The Genomic and Cellular Foundations of Animal Origins. Annu. Rev. Genet. 47:509–537.

Sathe S, Kaushik S, Lalremruata A, Aggarwal RK, Cavender JC, Nanjundiah V. 2010. Genetic Heterogeneity in Wild Isolates of Cellular Slime Mold Social Groups. Microb. Ecol. 60:137–148.

Schaap P, Winckler T, Nelson M, Alvarez-Curto E, Elgie B, Hagiwara H, Cavender J, Milano-Curto A, Rozen DE, Dingermann T, et al. 2006. Molecular Phylogeny and Evolution of Morphology in the Social Amoebas. Science 314:661–663.

Shrimali RK, Lobanov AV, Xu X-M, Rao M, Carlson BA, Mahadeo DC, Parent CA, Gladyshev VN, Hatfield DL. 2005. Selenocysteine tRNA identification in the model organisms Dictyostelium discoideum and Tetrahymena thermophila. Biochem. Biophys. Res. Commun. 329:147–151.

Singh B. 1947. Studies on Soil Acrasieae: 1. Distribution of species of Dictyostelium in soils of Great Britain and the effect of bacteria on their development. Microbiology 1:11–21.

Song J, Xu Q, Olsen R, Loomis WF, Shaulsky G, Kuspa A, Sucgang R. 2005. Comparing the Dictyostelium and Entamoeba Genomes Reveals an Ancient Split in the Conosa Lineage. Bourne P, editor. PLoS Comput. Biol. 1:e71.

Stanke M, Steinkamp R, Waack S, Morgenstern B. 2004. AUGUSTUS: a web server for gene finding in eukaryotes. Nucleic Acids Res. 32:W309–W312.

Stephenson SL, Landolt JC. 1992. Vertebrates as vectors of cellular slime moulds in temperate forests. Mycol. Res. 96:670–672.

Suthers HB. 1985. Ground-feeding migratory songbirds as cellular slime mold distribution vectors. Oecologia 65:526–530.

Thiel T, Michalek W, Varshney R, Graner A. 2003. Exploiting EST databases for the development and characterization of gene-derived SSR-markers in barley (Hordeum vulgare L.). Theor. Appl. Genet. 106:411–422.

Walker BJ, Abeel T, Shea T, Priest M, Abouelliel A, Sakthikumar S, Cuomo CA, Zeng Q, Wortman J, Young SK. 2014. Pilon: an integrated tool for comprehensive microbial variant detection and genome assembly improvement. PloS One 9:e112963.

Weis WI, Nelson WJ, Dickinson DJ. 2013. Evolution and Cell Physiology. 3. Using *Dictyostelium discoideum* to investigate mechanisms of epithelial polarity. Am. J. Physiol.-Cell Physiol. 305:C1091–C1095.

Wood DE, Lu J, Langmead B. 2019. Improved metagenomic analysis with Kraken 2. Genome Biol. 20:257.

Wu TD, Watanabe CK. 2005. GMAP: a genomic mapping and alignment program for mRNA and EST sequences. Bioinformatics 21:1859–1875.

Xiao Y, Stolzer M, Wasserman L, Durand D. 2025. Evolution of the Metazoan Protein Domain Toolkit Revealed by a Birth-Death-Gain Model. Available from: http://biorxiv.org/lookup/doi/10.1101/2025.07.03.659011

