## Supplementary Material for "A consensus genome sequence for the social amoeba *Dictyostelium giganteum*"

**A consensus genome sequence for the social amoeba *Dictyostelium giganteum* using Illumina reads.**

**Table S1: Transposable element families in *Dictyostelium giganteum* and their similarity to *D. discoideum.***

| **Subgroup** | **Class** | **Transposon** | **Avg. Identity to *D. discoideum* (%)** | **Range (%)** | **Avg. Length (bp)** | **Max Identity (%)** | **Hits** | **Highest Bitscore** |
| --- | --- | --- | --- | --- | --- | --- | --- | --- |
| **DNA transposons** | **DINOLT** | **DINOLT1** | 46.3 | 32.2–64.0 | 68 | 64.0 | 22 | 150.0 |
|  | **DRE** | **DRE** | 38.6 | 24.6–58.0 | 68 | 58.0 | 26 | 137.0 |
|  | **piggyBac** | **piggyBacA-1_DD** | 31.8 | 30.4–33.6 | 98 | 33.6 | 3 | 89.5 |
|  | **TDD** | **TDD3** | 50.7 | 28.6–100.0 | 54 | 100.0 | 54 | 180.0 |
|  |  | **TDD-4*** | 45.1 | 30.2–64.5 | 49 | 64.5 | 4 | 98.7 |
|  |  | **TDD-5** | 43.2 | 31.0–61.5 | 40 | 61.5 | 6 | 76.2 |
| **LTR transposons** | **DGLT-A** | **DGLT-A1_I** | 49.8 | 36.1–76.2 | 36 | 76.2 | 28 | 99.1 |
|  | **Gypsy** | **Gypsy-1-I_DD** | 43.6 | 28.1–62.5 | 36 | 62.5 | 7 | 27.6 |
|  | **DIRS** | **DIRS1** | 55.1 | 34.4–86.8 | 69 | 86.8 | 82 | 922.0 |
| **Non-LTR transposons** | **TRE** | **TRE3-A*** | 54.1 | 21.3–85.7 | 87 | 85.7 | 107 | 756.0 |
|  |  | **TRE3-B*** | 45.1 | 24.1–88.2 | 67 | 88.2 | 47 | 308.0 |
|  |  | **TRE3-C*** | 51.3 | 27.5–85.7 | 79 | 85.7 | 97 | 633.0 |

**Table S2: Comparative ncRNA Composition**

| **ncRNA Type** | ***D. giganteum*** | ***D. discoideum*** | ***D. firmibasis*** |
| --- | --- | --- | --- |
| **rRNA genes** | 16 | 15–18 | 15–18 |
| **tRNA genes** | 456 | ~420 | 379 |
| **Spliceosomal RNAs (U1–U6)** | 5 | 5 | 5 |
| **snoRNAs (DdR family)** | 8 | 8+ | 23 |
| **DUSE-associated structured RNAs** | 2 | 2 classes (I and II) | Not specified/ reported. |
| **Total distinct ncRNA families** | 55 | ~60 | ~55–60 |

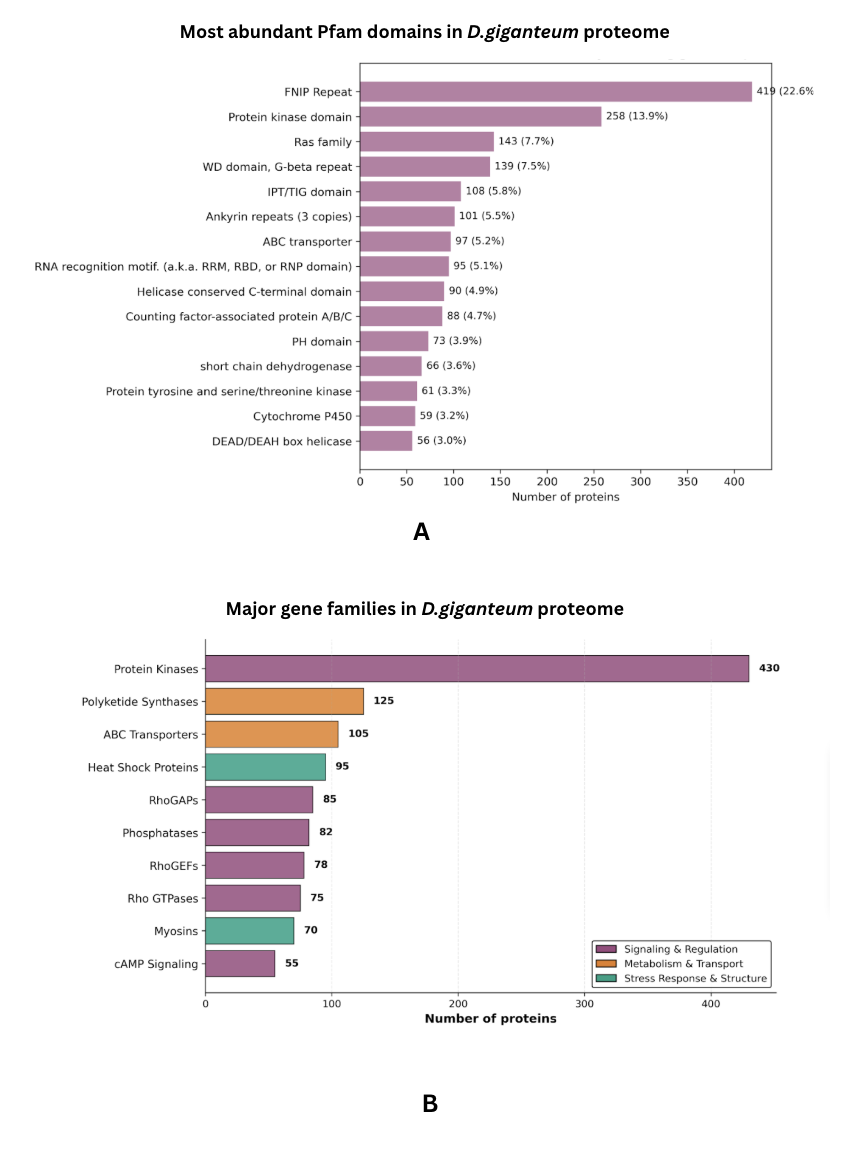

**Figure 1A: Most abundant Pfam domains in the *D. giganteum* proteome.**

The top 15 most abundant Pfam protein domains identified in the *Dictyostelium giganteum* proteome. Numbers indicate protein counts, with percentages showing the proportion of proteins containing each domain relative to the top 15 domains. FNIP repeats and protein kinase domains are the most prevalent, reflecting the importance of nutrient sensing and phosphorylation-based signalling in developmental regulation. The diversity of abundant domains highlights the regulatory complexity, metabolic versatility, and extensive post-transcriptional control mechanisms underlying multicellular development.

**Figure 2B. Major gene families in the *D. giganteum* proteome organized by functional category.**

Bar plot showing the abundance of selected expanded gene families in the *D. giganteum* predicted proteome. Protein kinases represent the largest category (430 proteins), followed by polyketide synthases (125) and ABC transporters (105). Additional expanded families include heat shock proteins (95), RhoGAPs (85), phosphatases (82), RhoGEFs (78), Rho GTPases (75), myosins (70), and components of the cAMP signaling pathway (55). Functional categories are color-coded as signaling and regulation (purple), metabolism and transport (orange), and stress response and structural proteins (green).

**Table S3:  Candidate horizontal gene transfers from Bacteria**

| **Pfam Domain** | **Function in Bacteria** | ***D. discoideum* Gene**  **(dictyBase ID)** | **Function in *D. discoideum*** | **Present in *D. giganteum*** | **Notes / Functional Inference in *D. giganteum*** |
| --- | --- | --- | --- | --- | --- |
| **Beta_elim_lyase** | Aromatic amino acid lyase | **DDB_G0281127** | Unknown | ✔ | Conserved lyase-like DUFs detected |
| **BioY** | Biotin transport/metabolism | **DDB_G0292424** | Unknown | ✔ | Biotin-binding/transport DUFs present |
| **Endotoxin_N** | δ-endotoxin-like domain | **DDB_G0289249** | Unknown | ✔ | Present; multiple toxin-like paralogs |
| **IPT/TIG** | Isopentenyl transferase | **DDB_G0277215** | Discadenine production | ✔ | Strongly expanded; signaling diversification |
| **IucA/IucC** | Siderophore synthesis | **DDB_G0294004** | Unknown | ✔ | Present; iron-scavenging functions retained |
| **OsmC** | Osmotic/oxidative stress | **DDB_G0268884** | Unknown | ✔ | Conserved stress-response proteins |
| **Peptidase S13/M15** | Dipeptidase / β-lactamase | **DDB_G0271902** | Carboxypeptidase | ✔ | Expanded hydrolase families |
| **PP_kinase** | Polyphosphate synthesis | **DDB_G0293524** | Polyphosphate synthesis | ✔ | Conserved across Dictyostelia |
| **TerD** | Tellurite resistance | **DDB_G0277501** | capA/B-related | ✔ | Present; detoxification role |
| **Thy1 (ThyX)** | Alternative thymidylate synthase | **DDB_G0280045** | Replaces canonical ThyA | ✔ | ThyX present; ThyA absent as in D. discoideum |
| **DUF885** | Unknown bacterial protein | **DDB_G0278355** | Unknown | ✔ | Highly expanded |
| **DUF1121** | Unknown | **DDB_G0277411** | Unknown | ✔ | Present |
| **DUF1289** | Unknown | **DDB_G0282477** | Unknown | ✔ | Present |
| **DUF1294** | Unknown | **DDB_G0285825** | Unknown | ✔ | Significantly expanded |
| **luc4/LucC** | Luciferase-like | **No dictyBase model ID provided** | Unknown | ✘ | Absent |
| **Cna_B** | Cell-adhesion repeat (Gram-positive) | **DDB_G0292696** | Colossin A | ✘ | Absent; lineage-specific loss |
| **Dyp_peroxidase** | Peroxidase | **DDB_G0273083** | Unknown | ✘ | Absent |
| **CnaB-embedded** | Adhesin-like repeats in large proteins | **Present in Colossin A (DDB_G0292696)** | Present in giant fusion proteins | ✘ | Absent; no CnaB-containing proteins detected |

**Comparison with *D. discoideum* and *D. firmibasis***

Comparative analysis of the nuclear genomes of *Dictyostelium giganteum*, *D. discoideum*, and *D. firmibasis* reveals broadly conserved genome organization with species-specific differences in size, gene architecture, and intron content (Figure 1). The *D.giganteum* genome spans 38.52 Mb, intermediate between *D. discoideum* (34 Mb) and *D.firmibasis* (31.5 Mb), and encodes 13,251 predicted protein-coding genes.

All three genomes exhibit pronounced AT-richness. The AT content of *D.giganteum* (75.76%) closely matches that of *D. discoideum* (77.57%) and *D. firmibasis* (76.01%), consistent with strong shared mutational biases across the genus. Gene density is comparable between *D.giganteum* and *D.discoideum*, whereas *D.firmibasis* displays lower gene density and longer average gene and intron lengths, suggesting relaxed structural compaction in the latter.

Despite these differences, mean protein lengths are highly conserved across species, indicating evolutionary constraint at the proteome level despite divergence in non-coding and intronic regions.

**Table S4: Comparative nuclear genome features of *Dictyostelium* species**

| **Feature** | ***D. giganteum*** | ***D. discoideum*** | ***D. firmibasis*** |
| --- | --- | --- | --- |
| **Genome size (Mb)** | 38.52 | 34.0 | 31.5 |
| **Protein-coding genes** | 13,251 | 13,541 | 10,564 |
| **Gene density (kb/gene)** | 2.41 | 2.50 | 2.98 |
| **Mean gene length (nt)** | 1,755.1 | 1,756.0 | 2,246.0 |
| **Introns per gene (spliced genes)** | 1.96 | 1.90 | 2.34 |
| **Mean intron length (nt)** | 129.5 | 146.0 | 183.1 |
| **Mean protein length (aa)** | 528.9 | 518.0 | 525.3 |
| **AT content (%)** | 76.76 | 77.57 | 76.01 |

### **Comparative mitochondrial genome organization**

The mitochondrial genome of *D.giganteum* is substantially smaller (50,452 bp) than those of *D. discoideum* (55,564 bp; 9.2% larger) and *D. firmibasis* (54,564 bp; 7.5% larger) (Figure 2). Despite reduced size, the *D.giganteum* mitochondrial genome exhibits the **highest AT content (75.7%)** among the three species, exceeding that of *D. discoideum* (72.6%) and *D. firmibasis* (71.1%).

Mitochondrial gene content shows moderate variation. *D.giganteum* encodes 36 protein-coding genes, fewer than *D.discoideum* (39) and *D.firmibasis* (38), primarily reflecting a reduced complement of oxidative phosphorylation (OXPHOS) genes. In contrast, the repertoire of mitochondrial ribosomal protein genes (rpl and rps) is fully conserved across all three species.

**Table S5: Comparative mitochondrial genome features of *Dictyostelium* species**

| **Feature** | ***D. giganteum*** | ***D. discoideum*** | ***D. firmibasis*** |
| --- | --- | --- | --- |
| **Mitochondrial genome size (bp)** | 50,452 | 55,564 | 54,564 |
| **Protein-coding genes** | 36 | 39 | 38 |
| **tRNA genes** | 16 | 15 | 17 |
| **16S rRNA (rnl, partial)** | 2 | 2 | 2 |
| **12S rRNA (rns)** | 1 | 1 | 1 |
| **AT content (%)** | 75.7 | 72.6 | 71.1 |
